# Conserved and divergent features of DNA methylation in embryonic stem cell-derived neurons

**DOI:** 10.1101/2020.01.08.898429

**Authors:** Sally Martin, Daniel Poppe, Nelly Olova, Conor O’Leary, Elena Ivanova, Jahnvi Pflueger, Jennifer Dechka, Rebecca K. Simmons, Helen M. Cooper, Wolf Reik, Ryan Lister, Ernst J. Wolvetang

## Abstract

DNA methylation functions in genome regulation and is implicated in neuronal maturation. Early post-natal accumulation of atypical non-CG methylation (mCH) occurs in neurons of mice and humans, but its precise function remains unknown. Here we investigate mCH deposition in neurons derived from mouse ES-cells *in vitro* and in cultured primary mouse neurons. We find that both acquire comparable levels of mCH over a similar period as *in vivo. In vitro* mCH deposition occurs concurrently with a transient increase in *Dnmt3a* expression, is preceded by expression of the post-mitotic neuronal marker *Rbfox3* (NeuN) and is enriched at the nuclear lamina. Despite these similarities, whole genome bisulfite sequencing reveals that mCH patterning in mESC-derived neurons partially differs from *in vivo*. mESC-derived neurons therefore represent a valuable model system for analyzing the mechanisms and functional consequences of correct and aberrantly deposited CG and non-CG methylation in neuronal maturation.

## Introduction

The unique epigenomic landscape of neurons is hypothesized to allow these post-mitotic cells to respond to diverse environmental stimuli during development and to modify gene transcription in response to activity, while retaining their cellular identity (Cortes-Mendoza et al., 2013, Day et al., 2013, Feng et al., 2010, Graff et al., 2012, Miller and Sweatt, 2007, Stroud et al., 2017). DNA methylation is thought to play an important role in imparting this simultaneous robustness and adaptability to neurons (Bayraktar and Kreutz, 2018b, Bayraktar and Kreutz, 2018a, Fasolino and Zhou, 2017). In most somatic cells, DNA methylation is largely restricted to cytosines in the context of CG dinucleotides (mCG). The methylation of CG sites is considered a relatively stable modification with a well-described function in gene silencing and imprinting (Bird, 2002). In contrast, in adult mammalian brains, other types of DNA methylation are found at high levels, including non-CG methylation (mCH, where H = A, T, or C (Guo et al., 2014, He and Ecker, 2015, Lister et al., 2013, Xie et al., 2012) and intermediates in the DNA demethylation pathway, particularly 5-hydroxymethylcytosine (5hmC) (Kriaucionis and Heintz, 2009, Lister et al., 2013, Mellen et al., 2012, Szulwach et al., 2011). In adult human and mouse neurons up to ∼50% of methylated cytosines in the genome occur in the mCH context, a level similar to mCG (Guo et al., 2014, Lister et al., 2013), and the majority of this exists in the mCA sequence context. While the precise roles of these modifications are not fully understood, the complex and diverse methylation profiles of adult neurons (Luo et al., 2017) suggest that DNA methylation plays an important role in the dynamic and adaptable regulation of gene expression in these cells. Studies in both mice and humans have shown that mCA is first observed in the brain shortly after birth and continues to accumulate during development to adulthood, after which the levels remain stable (Lister et al., 2013). This observation raises the exciting possibility that the generation of mCA may link early life experiences with neuron function later in life. The level of intragenic mCA in neurons inversely correlates with transcript abundance (Mo et al., 2015, Stroud et al., 2017, Xie et al., 2012), and in mice mCA deposition is negatively regulated by gene transcription (Stroud et al., 2017), suggesting that mCA functions as a part of a molecular system to modulate gene expression in response to synaptic activity, and to consolidate specificity of neuron subtypes.

DNA methylation is established and maintained by a family of conserved DNA (cytosine-5)-methyltransferases (DNMTs). Dnmt1 propagates existing methylation patterns at symmetrically opposed CG sites during cell division and is essential for the maintenance of methylation and chromosomal stability (Bayraktar and Kreutz, 2018a, Feng and Fan, 2009). Dnmt3a and Dnmt3b, on the other hand, catalyse the *de novo* methylation of cytosine, and the levels of Dnmt3a can be dynamically regulated to increase DNA methylation in the brain (Feng et al., 2005). The post-natal deposition of mCH is driven by a transient increase in the expression of Dnmt3a (Gabel et al., 2015, Guo et al., 2014, Lister et al., 2013, Luo et al., 2019, Stroud et al., 2017), and conditional deletion of Dnmt3a in Nestin-positive neuronal precursors during late gestation results in impaired motor activity (Nguyen et al., 2007). In contrast, deletion of Dnmt3a in excitatory neurons at early postnatal stages was reported to have no apparent major effect on brain development or function (Feng et al., 2010), suggesting that the developmental window during which mCH is deposited is critical (Lister et al., 2013). The importance of DNA methylation in governing correct neuronal function is exemplified by a range of developmental neurological disorders that result from mutations in proteins associated with DNA methylation in both the CG and CH context (Hamidi et al., 2015, Ip et al., 2018).

Defining the roles of mCG and mCH in neuron maturation and synaptic plasticity is of fundamental importance for understanding normal and abnormal brain development, thus a tractable and representative *in vitro* model system to further explore this process is highly desirable. We therefore investigated the levels, distribution and temporal dynamics of mCH during *in vitro* neuronal differentiation of human and mouse pluripotent stem cells. Deploying a range of cellular and genomic assays, we reveal similar sub-nuclear patterning, levels, and spatiotemporal dynamics of DNA methylome reconfiguration during the differentiation and maturation of mouse neurons *in vitro* and *in vivo*, but also differences that likely arise from the complex influence of the *in vivo* cellular environment on neuron differentiation and maturity.

## Results

### Immunocytochemical labelling for mCA accumulates in post-mitotic neurons and temporally correlates with DNMT3a expression

In vivo, mouse cortical neurons begin to acquire readily detectable levels of mCH around 2 weeks after birth, which continues to increase up to 6 weeks of age and remains high throughout adulthood (Lister et al., 2013). To assess whether this can be recapitulated in vitro, we used two independent approaches. First, we isolated primary cortical and hippocampal neurons from day 18 embryonic C57BL/6 mice (E18, average gestation 18.5 days) and cultured these for up to 14 days in vitro (DIV). We hypothesized that if mCH accumulation was due to cell intrinsic developmentally hardwired processes, 14DIV should correlate with the temporal acquisition of mCH in vivo (Figure 1A). In the second approach, we adapted an established differentiation protocol to generate mouse cortical neurons from embryonic stem cells (mESCs) (Bibel et al., 2007). We hypothesized that if this developmental model recapitulated neural development and neuronal maturation in vivo, mCH would occur within several weeks (Figure 1B). Two different mouse ESC lines, R1 (Nagy et al., 1993) and G4 (George et al., 2007), were differentiated as cell aggregates for 8 days in suspension, followed by dissociation and continued differentiation in adherent culture for up to 30 additional days to yield mixed cultures enriched in post-mitotic neurons (Supplementary Figure 1A). Both cell lines developed mature neurons within an equivalent time course and to a similar extent, as assessed by morphology and immunohistochemistry (R1 and G4), TEM analysis of synaptic depolarization (R1), and c-Fos mRNA and ICC analysis following depolarization (G4) (Supplementary Figure 1). To investigate the temporal acquisition, sub-nuclear localisation, and cell-type specificity of mCH in cultured neurons, we used an antibody raised against the mCA dinucleotide (anti-mCA) to analyse the primary- and mESC-derived neuronal cultures by immunocytochemistry. To identify neurons, cultures were co-labelled for NeuN/Rbfox3, a well-established marker of most postmitotic neuron subtypes, and beta3-tubulin (TUBB3), a pan-neuronal marker. Specificity of the mCA antibody was confirmed using a panel of competitive methylated oligonucleotides (Figure 1C – 1F, Supplementary Figure 2).

**Figure 1.**
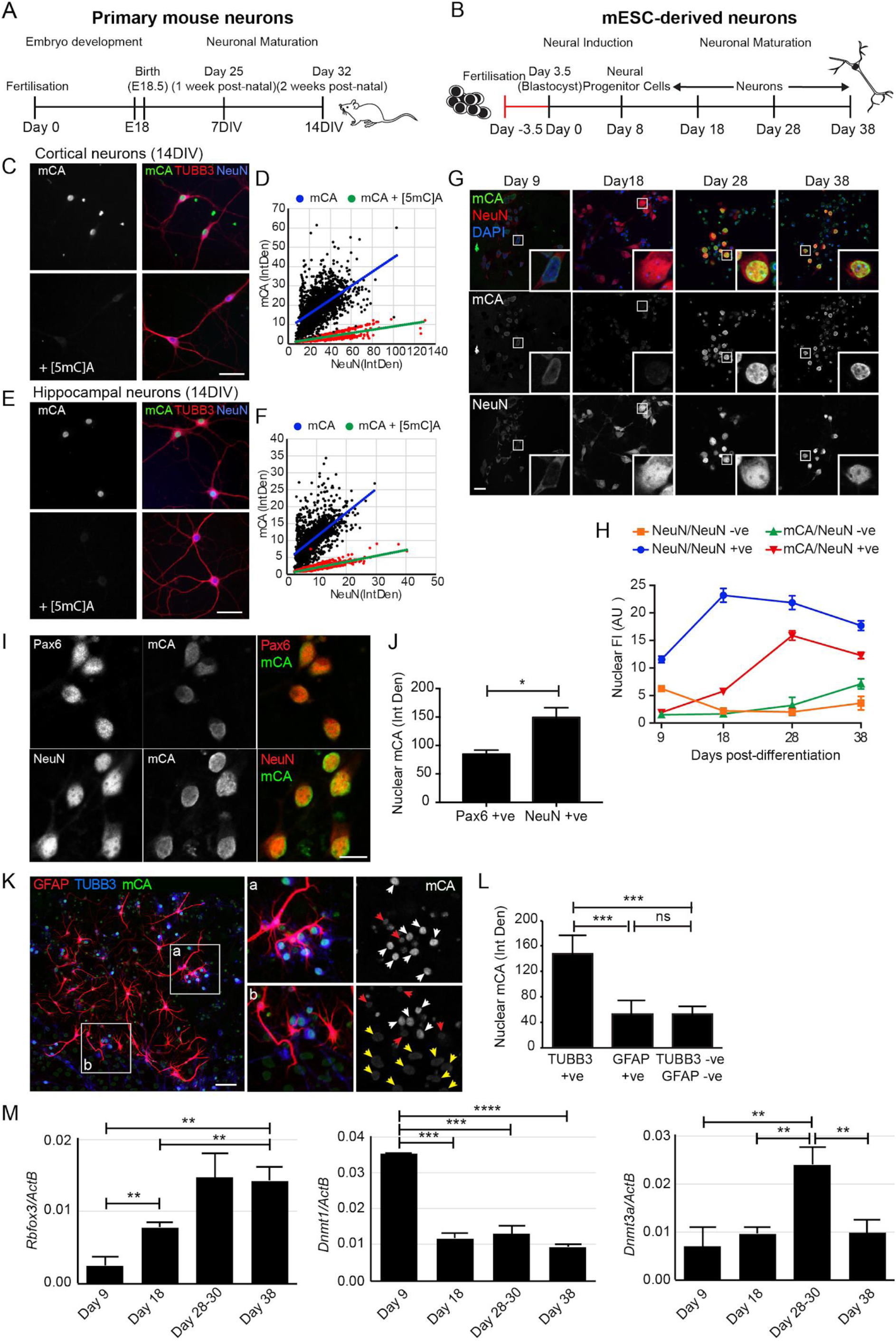
In vitro acquisition of DNA methylation in primary and ESC-derived mouse neurons. **(A, B)** Timeline schematic of neuron development in mouse cortex (A) and from mESCs in vitro (B). **(C-F)** 14DIV cortical (C, D) or hippocampal (E, F) neurons immunolabelled for NeuN, TUBB3 and mCA ± 2.5 µM [5mC]A. Scale bar = 50 µm. Analysis of the relative intensity of NeuN and mCA fluorescence in DAPI-masked nuclei from 14DIV cortical (D) or hippocampal (F) neurons. **(G, H)** mESC-derived neurons were fixed at different times during maturation and labelled for NeuN, mCA and DAPI. Scale bar = 20 µm. The nuclear fluorescence intensity (AU) of NeuN and mCA was determined. Results = mean ± SEM, for one representative differentiation. Similar labelling profiles were observed in 3 separate differentiations. **(I, J)** Neural progenitors (Day 9-10, Pax6+ve) and neurons (Day 20-40 NeuN+ve) were immunolabeled for mCA, and level of mCA nuclear fluorescence intensity quantified. Results = mean ± SD, n=3 separate differentiations for Pax6 and 2 differentiations for NeuN. Scale bar = 10 µm. **(K, L)** Later stage neural differentiations (Day 30-38) were immunolabeled for astrocytes (GFAP), neurons (TUBB3) and mCA. Nuclei were manually masked and the level of mCA measured in TUBB3+ve neuronal cells, GFAP+ve astrocytes, and GFAP-ve/TUBB3-ve cells of unknown identity. Quantitation shown was acquired from a single late-stage differentiation and labelling averaged across 5 random fields of view. Results are the mean ± SD. Arrows mark nuclei of astrocytes (red), neurons (white), and other cell types (yellow). Scale bar = 50 µm. **(M)** Gene expression of Dnmt1, Dnmt3a and Rbfox3 (NeuN) were determined by RT-qPCR relative to beta-actin. Results = mean ± SEM, n=3-4. For all experiments, statistical analysis was performed using a Student’s t-test. * p <0.01, ** p<0.05, *** p <0.001, **** p <0.0001.

To first determine whether *in vitro* cultured neurons could acquire non-CG methylation, 14DIV cortical and hippocampal primary neuronal cultures were immunolabelled for mCA, NeuN, and TUBB3 (Figure 1C, E). At this DIV, the majority of cells displayed a strong intranuclear labelling for mCA, in addition to labelling for both neuronal markers. Nuclear labeling for mCA was completely abrogated by the competitive [5mC]A oligonucleotide (Figure 1C-F, Supplementary Figure 2) confirming the specificity of the antibody. To quantify mCA labelling, cell nuclei were masked using DAPI fluorescence and the level of mCA and NeuN immunofluorescence in individual nuclei measured. The relative levels of mCA to NeuN were then determined (Figure 1D, 1F). Consistent with the enrichment of mCA in neurons, we found a strong positive correlation between the level of NeuN labelling and the level of mCA labelling in both hippocampal (r=0.62) and cortical (r=0.83) cells.

We next used the anti-mCA antibody to analyse the accumulation of mCA at different times up to 38 days during the differentiation and maturation of the mESC-derived neurons (Figure 1G, H). Neuronal cells expressed TUBB3 one day after attachment (day 9, Supplementary Figure 1E), and detectable NeuN labelling was observed within 3-6 days of attachment (differentiation day 11-14, Figure 1G), suggesting the rapid development of a post-mitotic phenotype. Interestingly, this temporally correlates with the initial identification of NeuN at E10.5 in the embryonic mouse brain (Mullen et al., 1992), suggesting that these developmental milestones are temporarily hardwired and can be recapitulated *in vitro*. Consistent with the early development of post-mitotic neurons, there was no further increase in the number of NeuN-positive neurons over the 4 weeks in adherent culture, although there was an obvious but variable increase in the number of non-neuronal cells within this time, including glial cells (see Figure 1K), as described previously (Bibel et al., 2004). Immunocytochemical (ICC)-based analysis of DNA methylation in NeuN-positive cells revealed an increase in the level of nuclear mCA labeling between days 18 and 28, which remained high to day 38 (Figure 1G, H). As the initial observation of mCA was significantly later than the initial observation of NeuN, a post-mitotic phenotype is likely a prerequisite for subsequent acquisition of mCA. Consistent with this, we found only minimal labelling for mCA in Pax6-positive neural progenitors differentiated for 9-10 days relative to ∼2-fold higher levels in NeuN-positive neurons differentiated for 28-38 days (Figure 1I, J). Similarly, we found minimal labelling for mCA in GFAP-positive glial cells and in additional unidentified cell types within the cultures that did not label for either neuronal or glial markers (Figure 1K, L).

Methylation of CH sites has been shown to be catalysed by Dnmt3a *in vivo* (Feng et al., 2005, Stroud et al., 2017). We therefore analysed the transcript abundance of *Rbfox3* (NeuN) and the DNA methyltransferases *Dnmt3a* and *Dnmt1* during differentiation by RT-qPCR (Figure 1M). Consistent with the results of the ICC, *Rbfox3* (NeuN) expression was found to significantly increase between days 9 and 18, reaching a plateau around day 28. In contrast, and in agreement with previous data from both primary neurons and mouse brain (Feng et al., 2005, Lister et al., 2013), *Dnmt3a* expression transiently increased between days 18 and 28 of differentiation and decreased again by 38 days. This transient increase in *Dnmt3a* transcript level temporally correlates with the accumulation of mCA labelling observed by ICC in the neurons (Figure 1G), strongly supporting the prediction that *Dnmt3a* catalyses CA methylation in the mESC-derived neurons. *Dnmt1* expression was initially high (Day 9) but rapidly decreased after cell attachment, consistent with a role in the maintenance of DNA methylation in the mitotic neural precursors. Collectively these data demonstrate that mCA deposition occurs specifically in post-mitotic neurons but not in non-neuronal cell types and correlates with DNMT3a expression.

### mCA associates with the nuclear lamina in neurons

During our ICC analysis of mCA labelling, we observed that this DNA modification was enriched near the nuclear periphery in our *in vitro* neuronal cultures. To determine whether this localisation was unique to mCA, ESC-derived neurons were co-labelled for total 5-methylcytosine (5mC) using an antibody predicted to label both mCG and mCH, and with the mCA-specific antibody, and the intranuclear distribution of the two compared (Figure 2A).

**Figure 2.**
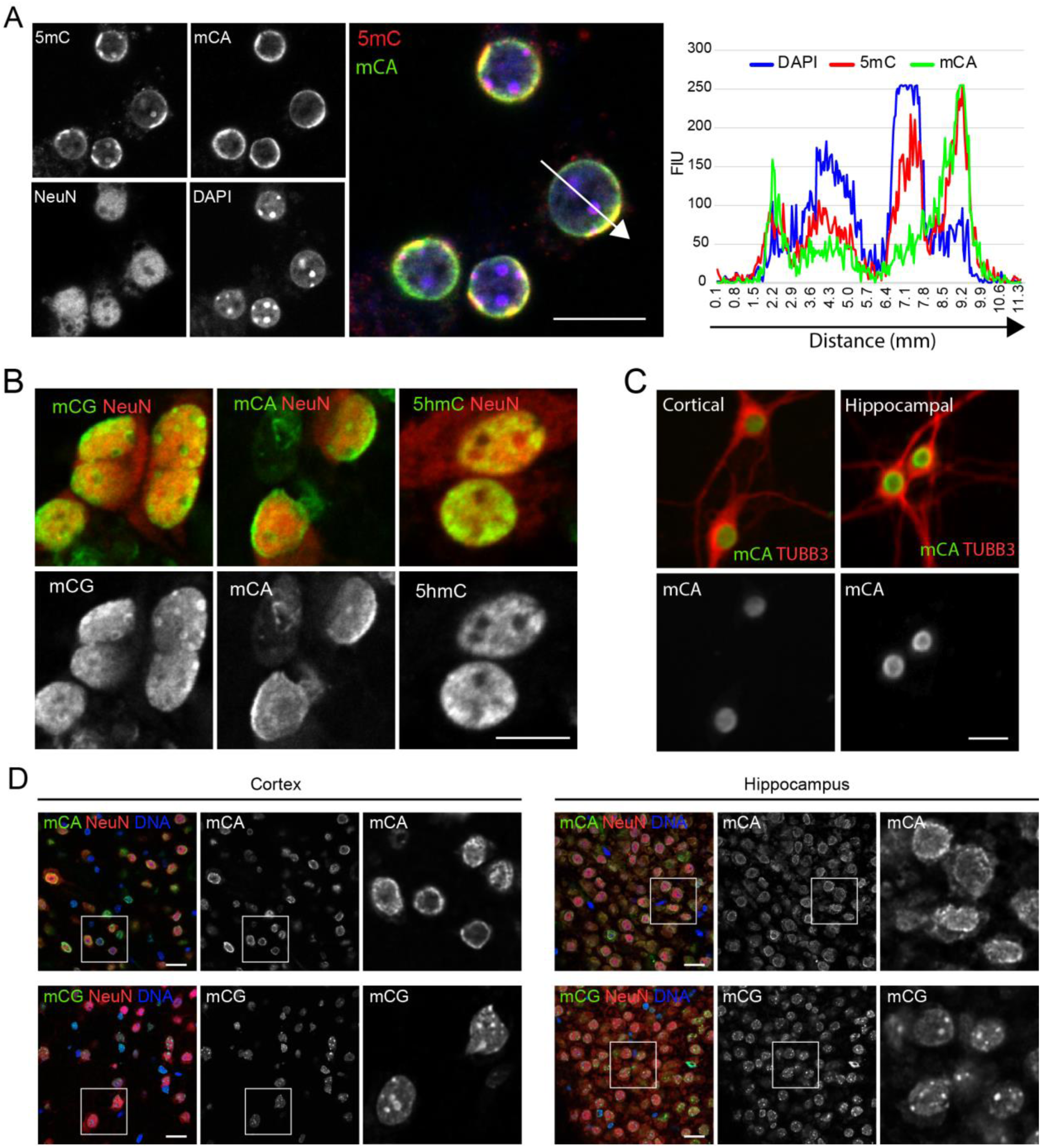
Intranuclear localisation of mCA in vivo, in primary mouse neurons and in mESC-derived neurons. **(A)** mESC-derived neurons were immunolabelled for total 5mC (mCG + mCH), mCA, and NeuN, and nuclei identified using DAPI. Scale bar = 10µm. **(B)** mESC-derived neurons were immunolabelled for NeuN and either mCG, mCA, or 5hmC. Scale bar = 10µm. **(C)** Primary mouse hippocampal and cortical neurons (14DIV) were labelled for mCA and TUBB3. Scale bar = 20µm. **(D)** Immunohistochemical analysis of DNA methylation in adult mouse hippocampus and cortex immunolabeled for either mCG or mCA, and NeuN, and nuclei identified using YOYO1. Scale bar = 20 µm.

Consistent with a more restricted intranuclear distribution of mCA, total 5mC labelling was observed to be more broadly distributed within the nucleus. While both marks showed a diffuse labeling, mCA was highly enriched at the nuclear periphery, whereas 5mC additionally strongly labelled intranuclear foci, which were devoid of mCA labelling. The intensely labelled 5mC foci were found to also stain strongly with DAPI (Figure 2A) suggesting regions of tightly packed heterochromatin that exclude mCA methylated DNA, in agreement with previous studies in mouse brain (Lister et al., 2013, Stroud et al., 2017).

To directly examine the distribution of DNA methylated in specific sequence contexts, neurons were labelled for either mCA or using an antibody specific for mCG (Figure 2B). In addition, we examined localisation of 5-hydroxymethylcytosine (5hmC) (Figure 2B), which is also highly enriched in neuronal genomic DNA (Kriaucionis and Heintz, 2009, Mellen et al., 2012, Szulwach et al., 2011). This labelling confirmed that the intense DAPI-stained foci observed in Figure 2A were enriched in mCG labelling. Again, mCA was enriched at the nuclear periphery, suggesting an association with the nuclear lamina. 5hmC was diffusely distributed throughout the nucleus, consistent with an enrichment in euchromatin (Chen et al., 2014).

To confirm that the distribution of mCA observed *in vitro* faithfully represented the intranuclear localisation *in vivo*, we analysed the localisation of mCA in primary neurons cultured for 14DIV (Figure 2C), and of mCA and mCG in immunohistochemical sections of adult mouse brain (Figure 2D). Both primary neurons and cortical or hippocampal brain sections showed an identical enrichment of mCA labelling at the nuclear periphery. Collectively these data show that DNA methylated in the CA context associates with the nuclear lamina in neurons.

### mESC-derived neuronal cultures acquire mCH to levels similar to those observed in vivo

Having shown that both mESC-derived neurons and cultured primary neurons acquired mCA labeling using immunocytochemistry, we next determined global levels of DNA methylation (CG and CH) by whole-genome bisulfite sequencing (WGBS).

In primary neurons, we found very low levels of mCH (<0.1%) at E18 in cells isolated from either the hippocampus or the cortex, consistent with previous *in vivo* data (Lister et al., 2013) and E12.5 mouse NPCs cultured *in vitro* (Luo et al., 2019). Following 14DIV, this level increased to 0.70% and 0.79% in primary cortical and hippocampal neurons, respectively (Figure 3D), confirming that the mechanisms underpinning accumulation of mCH are conserved in *in vitro* cultures. This is in agreement with the *in vitro* differentiation of E12.5-derived mouse NPCs, which also show an increase in mCH levels over several weeks of culture, reaching a maximum of 0.35% mCH/CH after 21 days (Luo et al., 2019). Analysis of mCH context confirmed that methylation at CA sites was the most abundant modification, although smaller increases in methylation in the CT and CC contexts were also observed (Figure 3E-G). Comparison of the methylation levels at 14DIV to those of 2 week old mouse prefrontal cortex (PFC) (Lister et al., 2013) showed that these were similar for all three mCH subtypes. In contrast, the levels of mCG were slightly higher *in vitro* than *in vivo* (Figure 3C).

**Figure 3.**
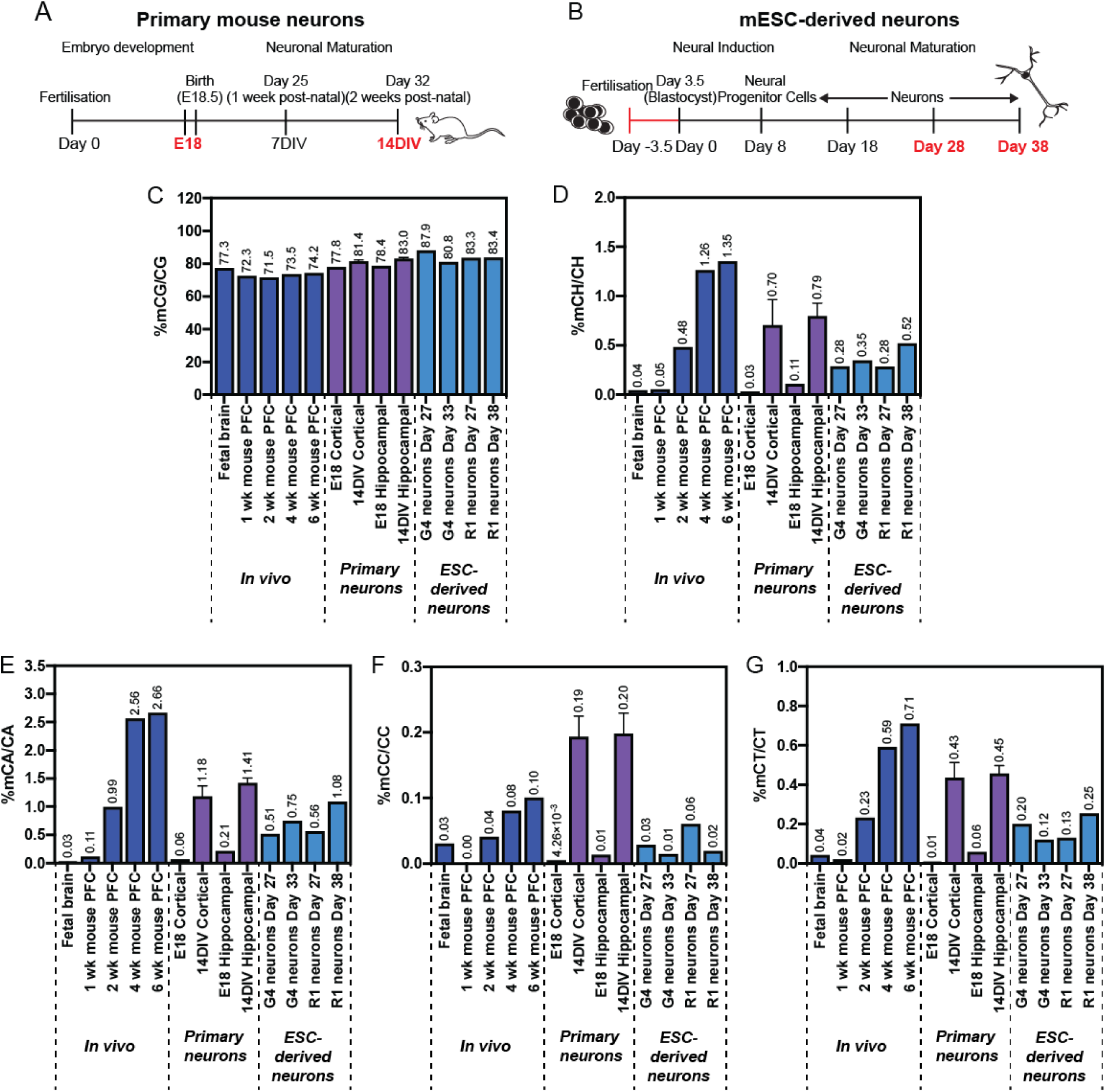
WGBS analysis of DNA methylation in primary mouse neurons and mESC-derived neurons. **(A, B)** Timeline schematic of neuron development in mouse cortex (A) and from mESCs in vitro (B). Time points analysed by WGBS are highlighted red. **(C-G)** Global levels of DNA methylation in mouse brain ((Lister et al., 2013), dark blue bars), primary mouse cortical or hippocampal neurons (E18 and 14DIV, purple bars) and mESC-derived neuronal cultures (R1 and G4 cell lines, light blue bars). Values represent the weighted methylation levels: the fraction of all WGBS base calls that were C at cytosine positions in the genome (for each context separately). Results = single samples, except 14DIV where n=2, mean ± SD.

Analysis of global DNA methylation levels by WGBS was then undertaken on the mESC-derived neuronal cultures between days 27 and 38 post-differentiation, which are temporally comparable to approximately 1 week and 3 weeks post-natal *in vivo* development, respectively (Figure 3A, B). Consistent with the primary neuron analysis, both the G4- and R1-derived neuronal cultures acquired high levels of mCH within 27 days, consisting predominantly of mCA. The global level of mCH and mCA was very similar between the two cell lines and showed a time-dependent increase up to 38 days. Global levels of mCA and mCH at 38 days (equivalent to ∼3 weeks post-natal *in vivo*) closely mirrored those of the prefrontal cortex of 2 week old mice (Figure 3D, E), although the levels at 27 days (∼1 week post-natal *in vivo*) were ∼5-fold higher than in 1 week old mouse PFC. Whether this reflects an earlier deposition of mCH *in vitro* or is a result of differing proportions of neuronal and non-neuronal cells in the two sample types is not known. Smaller increases in the level of mCT and mCC were also observed *in vitro*, as well as an increase in the level of mCG. Our data indicate that *in vitro* neuronal differentiation of mESCs recapitulates overall *in vivo* levels of mCH and mCA, which is distinct from prior report of low levels of mCH and mCA in iN cells (Luo et al., 2019). Thus, we conclude that acquisition of mCH and mCA is largely a neuro-developmentally hardwired process.

### Global mCH and mCG levels in mESC-derived neurons reveals hypermethylation relative to *in vivo* adult neurons

As mESC-derived neuronal cultures contain multiple cell types (Figure 1) fluorescence-activated nuclear sorting (FANS) was used to analyse mCG and mCH levels in specific populations of cells (Figure 4A, B). To analyse the developmental timeline of mESC-derived neurons we sorted Nanog-positive nuclei from mESCs, Pax6-positive nuclei from mESC-derived neuronal progenitor cells, NeuN-positive nuclei from cells cultured for 30 days, and NeuN-positive and NeuN-negative nuclei from cells cultured for 38 days (Figure 4A, see also Figure 1A, B). To investigate a possible temporal regulation of methylation patterns we also isolated NeuN-positive cells from various human ESC-derived neuronal cultures, including both 2-D cortical differentiation (Reinhardt et al., 2013) and 3-D cerebral organoids (Lancaster and Knoblich, 2014) (Figure 4B). We then analysed by WGBS the levels of mCG and mCH in these different nuclear populations, representing different cell types within the various neuronal differentiation timelines (Figure 4A and 4B).

**Figure 4.**
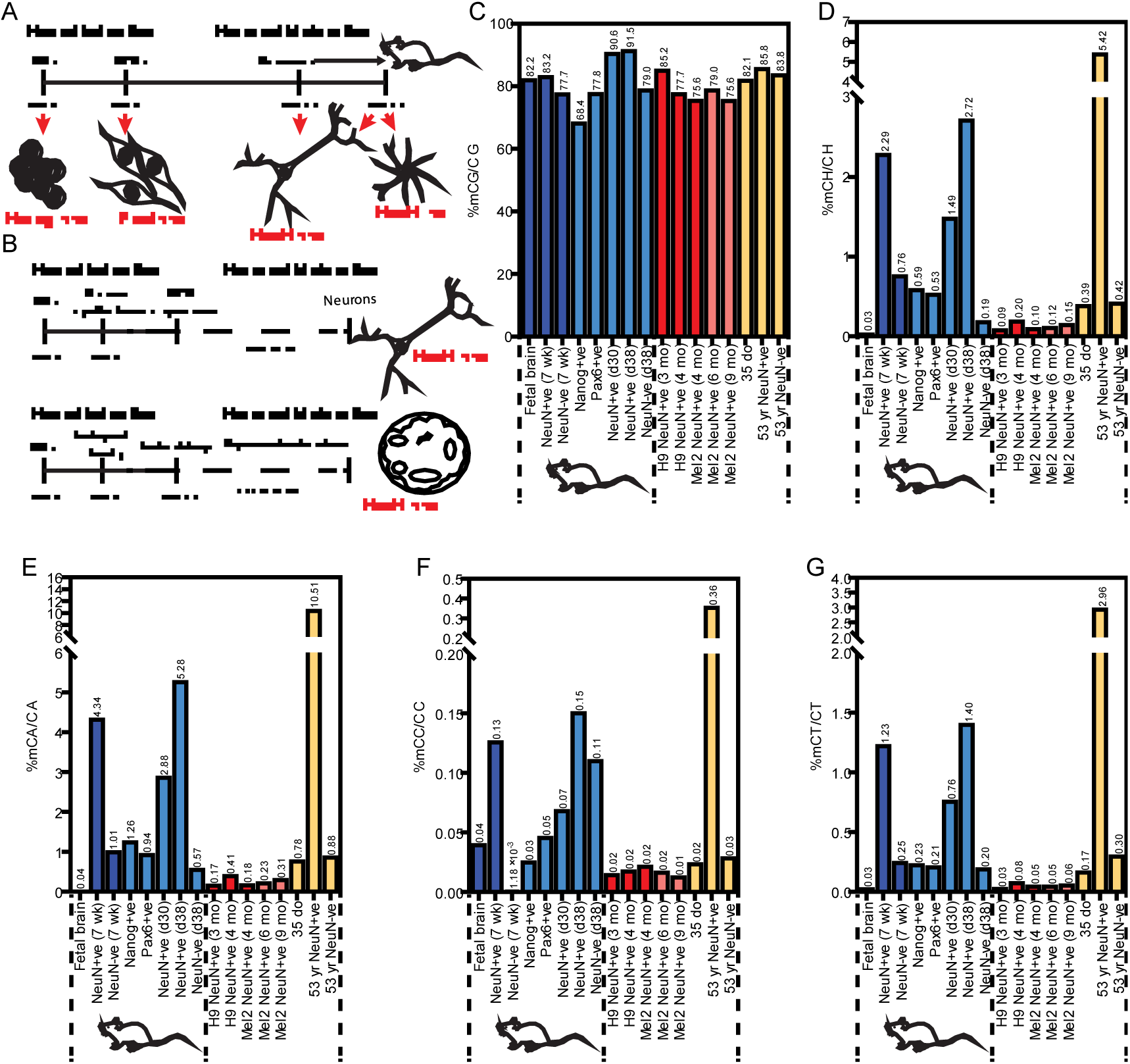
DNA methylation levels in fluorescence-activated sorted nuclei. **(A, B)** Schematics showing time points during the mESC and hESC differentiation to neurons at which nuclei were isolated. **(A)** Samples from the mESC differentiation included Nanog +ve mESCs, Pax6 +ve neural progenitors, day 30 and 38 NeuN +ve mouse neurons and day 38 NeuN-ve cells. **(B)** In the human ESC differentiation, NeuN +ve nuclei were isolated following 12-16 weeks of 2-D culture, or 6-9months of 3-D culture. **(C-G)** Level of DNA methylation in isolated nuclear populations. Light blue (mouse), red (human 2-D) and pink (human 3-D) bars show samples generated in this study. Dark blue (mouse) and yellow (human) bars show previously published levels of DNA methylation (Lister et al., 2013).

During the mouse ESC differentiation process, global CG methylation levels were observed to progressively increase, with the highest increment observed during the maturation of Pax6-positive neural progenitors to 30-day old neuronal nuclei, corresponding to the period during which maximal *Dnmt3a* transcript abundance was observed in bulk cultures (Figure 1M). The observed levels of mCG in NeuN-positive nuclei, at both 30 days and 38 days, was substantially higher than that reported in NeuN-positive nuclei isolated from mouse PFC, and was higher than the equivalent NeuN-negative cells in the cultures, suggesting specific hypermethylation of neuronal CG occurs *in vitro*. Interestingly, hypermethylation of CG was not observed in the human cultures, where we observed generally lower global levels of mCG than *in vivo* (for four out of five cultures).

We next analysed the levels of mCH in the various nuclei populations. In the mouse samples, the level of mCH was found to increase substantially during the transition from Pax6-positive NPCs to 30 day old NeuN-positive neurons, and again between day 30 and day 38 of neuron maturation (Figure 4D). The levels attained by day 38 of culture exceeded that of 7 week old mouse prefrontal cortex (approximately 65 days total development from the blastocyst stage), suggesting that for mCH, as for mCG, hypermethylation of the *in vitro*-derived neurons was occurring. This pattern of methylation was recapitulated for the individual analyses of mCA, mCC and mCT (Figure 4E-F). For mCA, we found a level of 1.25% (mCA/CA) in Nanog-positive mESC-derived nuclei, consistent with published studies in mice (Arand et al., 2012, Ramsahoye et al., 2000) and humans (Liao et al., 2015, Ziller et al., 2011). This level was found to decrease in Pax6-positive neural progenitors, and subsequently increased to 2.9% (day 30) and 5.3% (day 38) in NeuN-positive nuclei. As with mCG and mCH, at day 38 this level was higher than that reported in NeuN-positive nuclei isolated from mouse brain, again suggesting hypermethylation of CA. Consistent with the ICC results and published *in vivo* data (Lister et al., 2013), NeuN-negative cells within the 38-day old neuronal cultures contained low levels of mCA. The levels of mCT in the day 38 ESC-derived neurons reached levels similar to those in the adult mouse brain, while only negligible levels of mCC were detected in any cell type. Together these data demonstrate that mESC-derived neurons acquire non-CG methylation levels similar to *in vivo* levels.

In contrast to mouse ESC-derived neurons, only negligible levels of non-CG methylation were observed in any of the human ESC-derived neuronal populations, suggesting that even temporally extended human neuronal cultures are unable to mature sufficiently to acquire mCH. No difference in mCH was detected between shorter, 2-D neuronal cultures, and aged cerebral organoids, suggesting that the culture conditions alone do not promote the acquisition of mCH.

### Genome wide distribution of mCH and mCG DNA methylation between *in vivo* and mESC-derived neurons indicates regional hypermethylation

To establish the degree to which DNA methylation patterns in the ESC-derived neurons recapitulated those of *in vivo* neurons, we generated base resolution methylomes by WGBS and assessed the regional distribution of mCG (Figure 5) and mCH (Figure 6). We then compared this to previously published datasets from 7-week old mouse PFC glial cells (glia) or NeuN-positive neurons (*in vivo* neurons), and fetal mouse frontal cortex (fetal)(Lister et al., 2013), in order to identify potential differences between *in vivo* and *in vitro* neurons (Figures 5 and 6) and the similarities (Figure 7).

**Figure 5.**
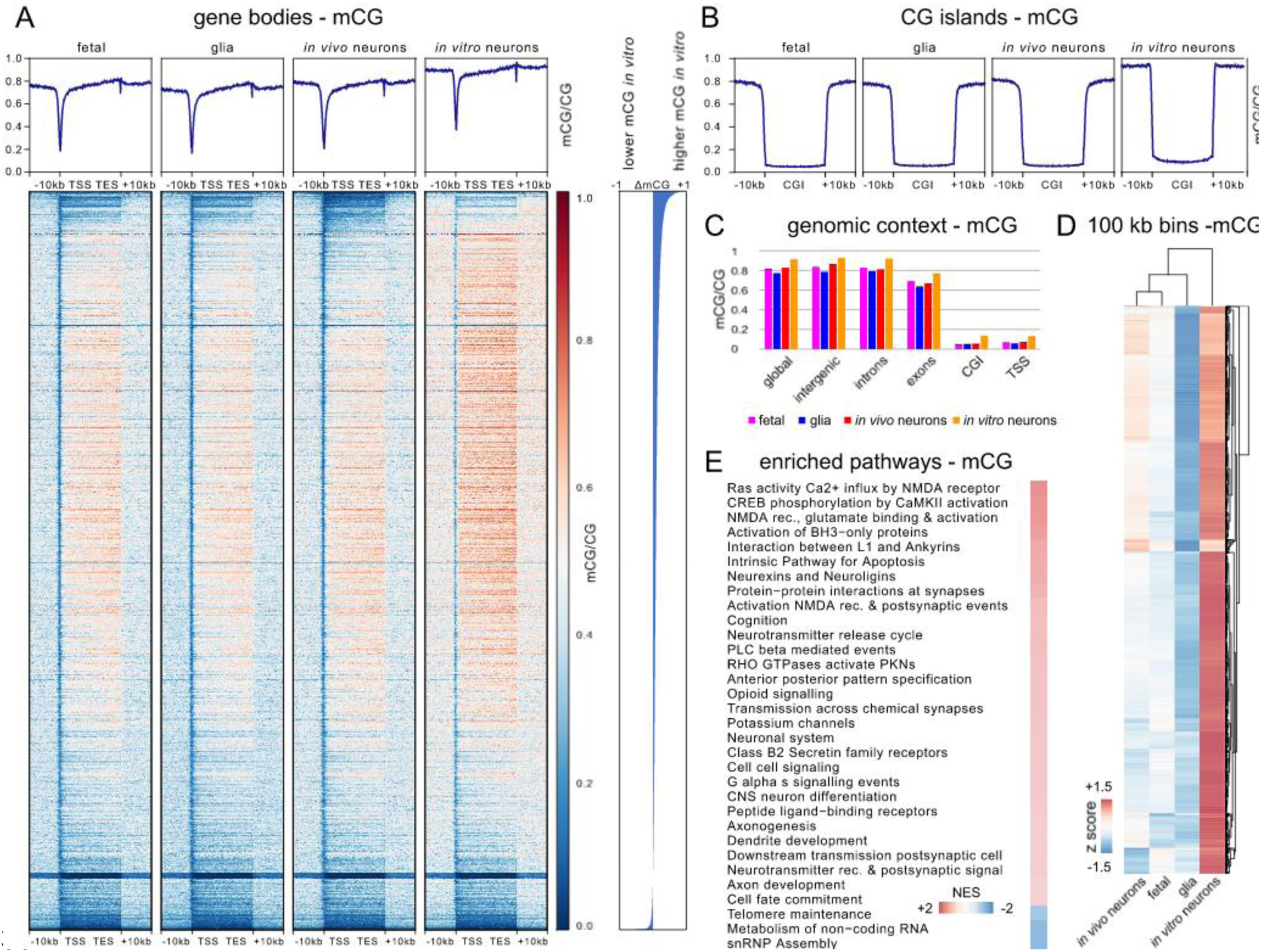
Global mCG properties of in vivo vs in vitro generated neurons. mCG characteristics for d38 mESC-derived neurons (in vitro neurons), 7-week old mouse prefrontal cortex neurons (in vivo neurons), NeuN-negative cells from 7-week old mouse prefrontal cortex (glia), and fetal mouse frontal cortex (fetal). **(A)** Weighted methylation levels (mCG/CG) for all genes and 10 kb flanking regions shown at top. Heatmap shows mCG of genes and flanking regions sorted by difference in gene body mCG/CG between mESC-derived neurons and in vivo adult mouse PFC neurons, as indicated on the right of the heatmap. **(B)** Weighted CG methylation level throughout CpG islands (CGIs) for all CGIs and 10 kb flanking regions. **(C)** Weighted methylation levels (mCG/CG) for the whole genome, intergenic regions, introns, exons, CGIs, and 500 bp flanking transcription start sites (TSS). **(D)** Hierarchical clustering based on Spearman correlation of mCG levels in all 10 kb bins of the genome. **(E)** Enriched pathways after pre-ranked gene set enrichment analysis based on differences in gene body mCG/CG between in vivo and in vitro neurons. Shown are top pathways based on enrichment scores (NES) for genes with higher mCG/CG in vitro (positive NES score) and lower mCG/CG in vitro (negative NES score).

**Figure 6.**
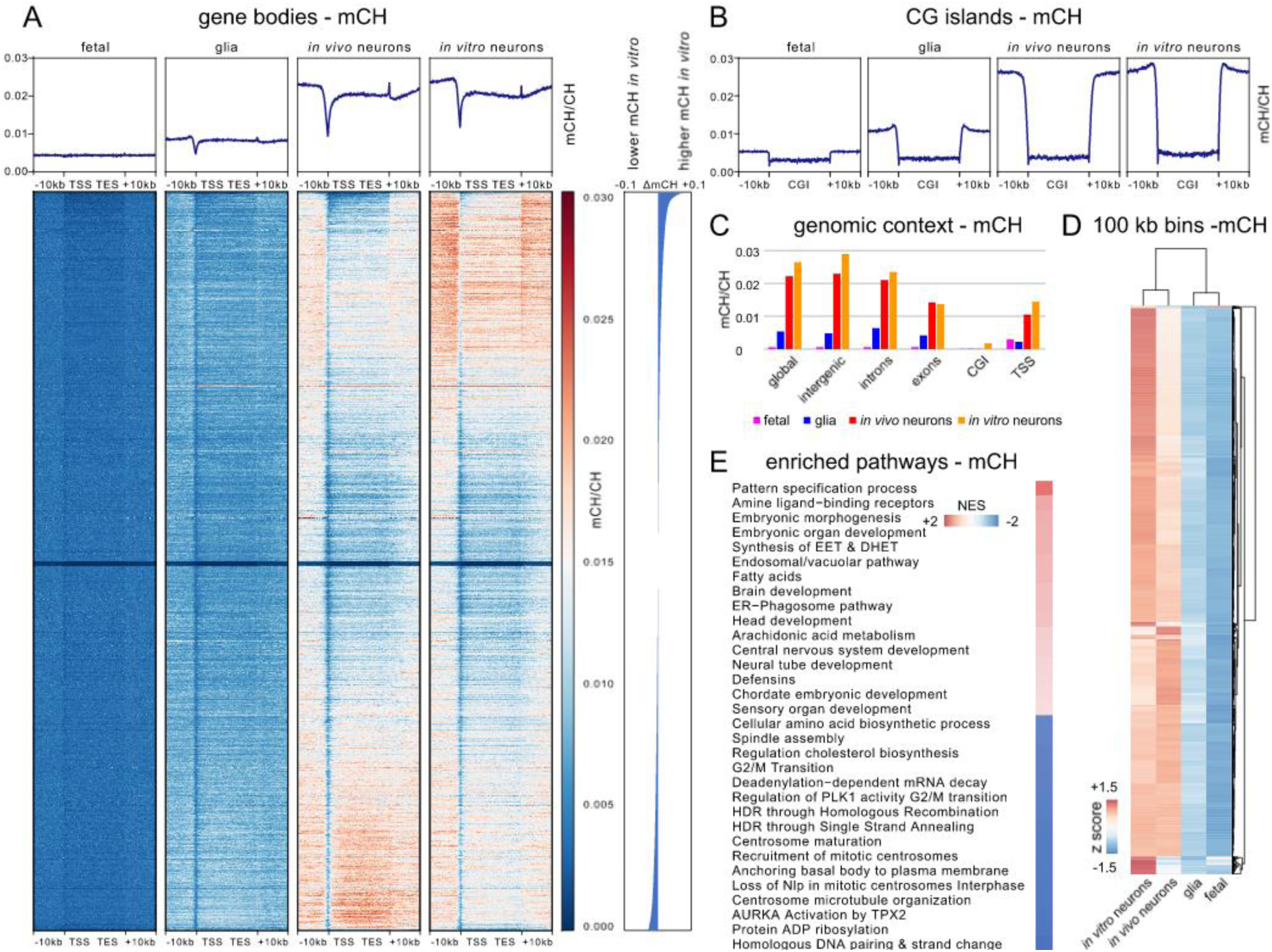
Global mCH properties of in vivo vs in vitro generated neurons. mCH characteristics for d38 mESC-derived neurons (in vitro neurons), 7-week old mouse prefrontal cortex neurons (in vivo neurons), NeuN-negative cells from 7-week old mouse prefrontal cortex (glia), and fetal mouse frontal cortex (fetal). **(A)** Weighted methylation levels (mCH/CH) for all genes and 10 kb flanking regions shown at top. Heatmap shows mCH of genes and flanking regions sorted by difference in gene body mCH/CH between mESC-derived neurons and in vivo adult mouse PFC neurons, as indicated on the right of the heatmap. **(B)** Weighted CH methylation level throughout CpG islands (CGIs) for all CGIs and 10 kb flanking regions. **(C)** Weighted methylation levels (mCH/CH) for the whole genome, intergenic regions, introns, exons, CGIs, and 500 bp flanking TSSs. **(D)** Hierarchical clustering based on Spearman correlation of mCH levels in all 10 kb bins of the genome. **(E)** Enriched pathways after pre-ranked gene set enrichment analysis based on differences in gene body mCH between in vivo and in vitro neurons. Shown are top pathways based on enrichment scores (NES) for genes with higher mCH/CH in vitro (positive NES score) and lower mCH/CH in vitro (negative NES score).

**Figure 7.**
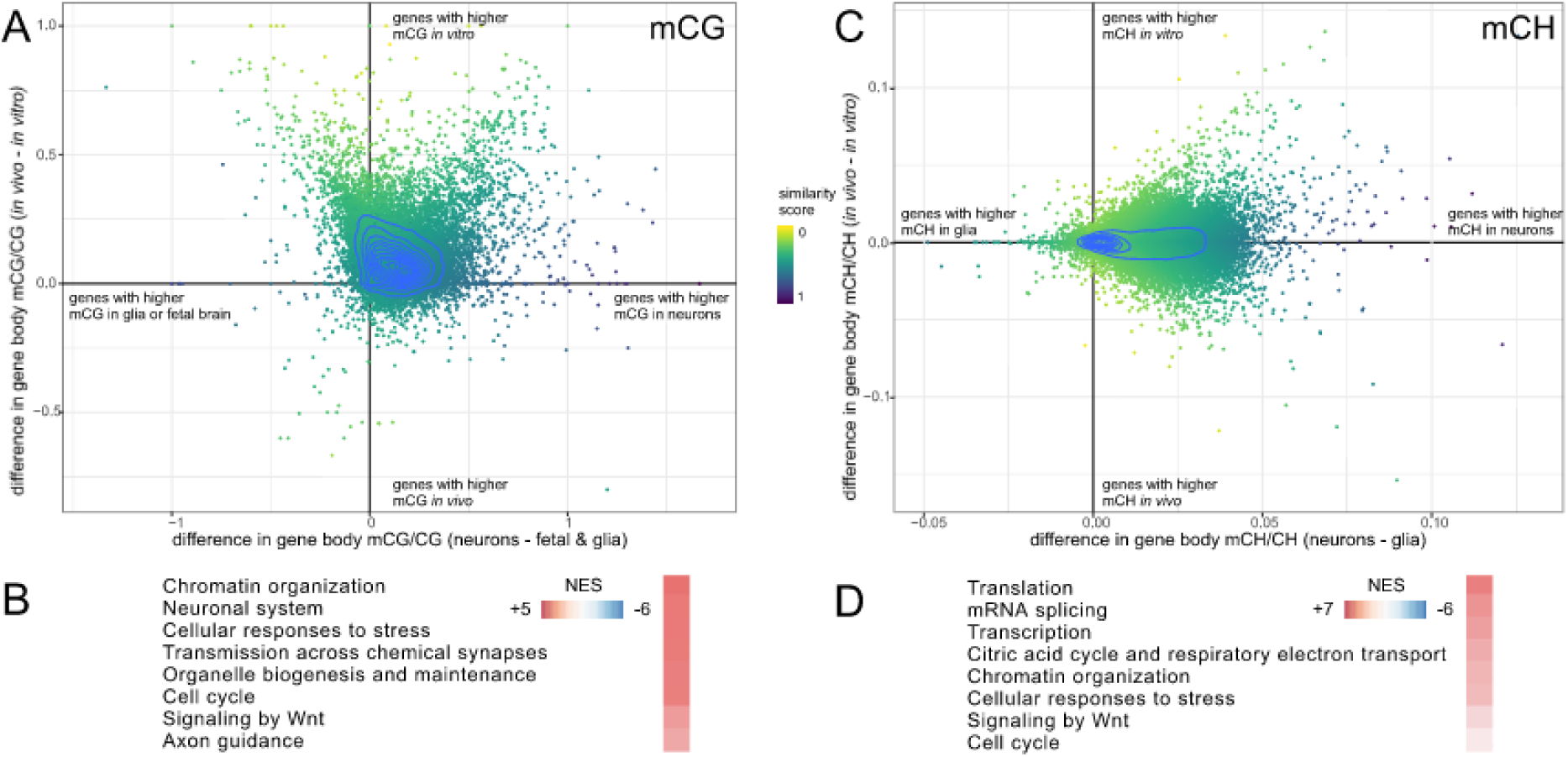
Enrichment for genes with similar methylation patterns in in vitro neurons and in vivo neurons. **(A)** Differences in gene body mCG level between neurons and fetal brain as well as glia (x-axis), and differences in gene body mCG level between both neuronal populations (y-axis), for all 24,049 genes with WGBS coverage. Dots are coloured by the similarity score that was used for gene set enrichment. Blue contour plot is based on gene density. **(B)** Selection of top enriched pathways from GSEA using similarity scoring from mCG information. **(C)** Differences in gene body mCH level between neurons and glia (x axis), and differences in gene body mCH level between both neuronal populations (y-axis), for all 24,049 genes with WGBS coverage. **(D)** Selection of top enriched pathways from GSEA using similarity scoring from mCH information. Full details for the top 20 similarity-enriched pathways are listed in Supplementary Figure 11.

Analysis of average mCG levels across all gene bodies and associated 10 kb flanking regions (Figure 5A) showed that these were generally similar between fetal, glial and *in vivo* adult neurons, but exhibited a generalised increase in ESC-derived neurons (designated as *in vitro* neurons) (Figures 5A, top plots), consistent with the observed global mCG levels (Figure 4). Analysis of mCH levels (Figure 6A) showed that these were very similar between *in vivo* and *in vitro* neurons, although there was a small localised increase around transcription start sites *in vitro*, and they were much higher than either the glial or fetal samples (Figure 6A, top plot). Genes were subsequently ordered by difference in mean gene body methylation level relative to *in vivo* neurons (Figure 5A and 6A, lower heatmaps). In this analysis, we observed that 38.7% of genes were mCG hypermethylated (gene body ΔmCG > 0.1) in *in vitro* neurons compared to *in vivo* neurons, while only 0.6% of genes showed mCG hypomethylation (ΔmCG > 0.1). In the CH context the ratio was more balanced, with 16.0% of genes hypermethylated with ΔmCH > 0.01, and 12.2% of genes hypomethylated with ΔmCH > 0.01. Thus, while global and average mCG and mCH gene methylation levels are similar between *in vitro* and *in vivo* neurons, differences in the level of gene body DNA methylation are evident. As a recent study had described DNA methylation patterns in directly reprogrammed mouse neurons (iN cells) (Luo et al., 2019), we also compared our gene body methylation data to the iN cell data (Supplementary Figures 3 - 4). This analysis showed that the hypermethylation of mCG observed in mESC-derived neurons was unique to this developmental model, as both the levels and the patterns observed in iN cells were very similar to those of the *in vivo* neurons (Supplementary Figure 3). For mCH, the overall level of gene body methylation in iN cells was very low (mCH/CH) and lacked the localised spike in methylation observed at TESs *in vivo* and in ESC-derived neurons (Supplementary Figure 4). To directly compare methylation patterns and compensate for the lower overall mCH levels, gene body methylation levels were normalised to the average global methylation level within individual neuron populations. For mCG, despite the slightly lower methylation levels in the iN cells, when methylation patterns were normalised, a higher degree of similarity was observed to the mESC-derived neurons, suggesting that neither *in vitro* neuron population attains genic mCG patterning identical to *in vivo* neurons (Supplementary Figure 3). For mCH, more complex differences were observed, with both iN cells and ESC-derived neurons showing regional variation in the degree of similarity to *in vivo* neurons (Supplementary Figure 4).

To investigate the relationship between mCG and mCH within gene bodies in the ESC-derived neurons, we assessed mCH patterns over all genes ordered by gene body mCG difference (*in vitro* neuron mCG - *in vivo* neuron mCG; Supplementary Figure 5), and gene body mCG patterns over all genes ordered by gene body mCH difference (*in vitro* neuron mCH - *in vivo* neuron mCH; Supplementary Figure 7). These analyses revealed that the patterns of DNA methylation levels observed were different for each context (Figure 5A and 6A). Indeed a very low Pearson correlation of differences in methylation for both contexts between ESC-derived neurons and in vivo neurons (r = 0.0688) suggests that independent processes define DNA methylation levels for each context, and that individual genes are methylated to a different degree by mCG and mCH, compared to the global average levels. Plotting the total number of calls at cytosine reference positions within gene bodies in the same order confirmed that the observed differences are not a result of different sequencing coverage between samples (Supplementary Figures 6 and 8).

In mESC-derived neurons, a marginal and consistent increase in mCG was observed within and flanking CpG islands (CGIs), regions that are normally depleted of methylation (Schubeler, 2015) (Figure 5B). Similarly, there was a slight increase in mCH methylation within CGIs. However, there was a pronounced and localised increase in the level of mCH in the regions immediately flanking CGIs (Figure 6B), suggesting that methylation of CG and CH sites in these regions are regulated differently. In order to assess whether particular genomic features exhibited different methylation levels in mESC-derived neurons, we measured the weighted methylation levels (mCG/CG or mCH/CH) in intergenic regions, introns, exons, and 500 bp upstream and downstream of transcription start sites (Figures 5C and 6C). This revealed genome-wide CG hypermethylation in mESC-derived neurons (∼6-12% higher than in *in vivo* neurons, absolute methylation level difference), suggesting a generalised dysregulation of methylation level, while increased mCH was observed in all genomic features except exons. The reason for the specific exclusion of exons from hypermethylation in the mCH context is unknown but suggests that specialised mechanisms regulating exonal mCH levels are conserved *in vitro*.

Next, we assessed regional correlation in mCG and mCH levels in 100 kb bins of the whole genome (excluding chromosomes X and Y) between mESC-derived neurons, fetal frontal cortex, and 7-week old mouse PFC neurons and glia (Figure 5D and 6D). For mCG, fetal brain and adult neurons were the most similar, with adult glia joining at the next node, while mESC-derived neurons formed their own branch. The low correlation between neuronal samples is likely due to the overall higher methylation of CG in ESC-derived neurons. For mCH, there was a high similarity between *in vitro-* and *in vivo*-derived neuronal datasets, while glia were similar to the fetal sample, consistent with the neuron-specific accumulation of mCH. This clustering based on bins also showed that differences in methylation between neurons were not evenly distributed throughout the genome but show regional variability.

Finally, we generated ranked lists of genes based on either similarity or difference in genic methylation between neuronal populations and performed gene set enrichment analysis (GSEA) to examine correlation with biologically relevant pathways. We composed a pathway package by combining gene ontology as well as reactome pathway datasets and performed GSEA on both gene lists. Initially, we performed GSEA based on differences in methylation levels (Figure 5E and 6E). For mCG, top pathways with higher gene body methylation *in vitro* represented neuronal activity and synapse formation. Due to the generalised hypermethylation in the CG context, relatively few pathways were enriched for genes with lower methylation values (at p<0.05, Figure 5E), none of which directly related to neurons. Full details of the top 50 hyper- and hypomethylated pathways for mCG context are listed in Supplementary Figure 9. For mCH, top pathways enriched for genes hypermethylated in *in vitro* neurons included morphogenesis and development, including brain and central nervous system development. However, unlike mCG, differences in neuronal activity and synapse formation were not detected. Genes hypomethylated in the mCH context in *in vitro* neurons included pathways for cell cycle, cell division and DNA repair, but again, no pathways relating directly to neurons were identified (Figure 6E). Full details for the top 50 of hyper- and hypomethylated pathways for mCH context are listed in Supplementary Figure 10. To simultaneously identify gene sets that share similar methylation states in both *in vitro* and *in vivo* neuron populations and discriminate them against fetal or glial cells, we applied GSEA on genes ranked by a combination of similarity between the mESC-derived *in vitro* neurons and *in vivo* neurons, and dissimilarity to non-neuronal cell types (glia and fetal brain cells, Figure 7). This analysis was performed for mCG and mCH independently and resulted in an enrichment for pathways linked to genes that have an equivalent methylation state in both of the neuronal samples. It is important to note that this analysis enriches for similarity irrespective of overall methylation levels, whether high or low. When considering mCG, the most represented pathways belonged to two major groups: neuronal function or cell cycle. While neuronal terms are expected to be shared between neurons, the enrichment of the latter group is likely due to its associated genes having a different methylation state in post-mitotic neurons, compared to actively proliferating cells. Interestingly for mCH, with the exception of Wnt signalling, the most highly enriched pathways did not relate directly to neurons, but included pathways related to chromatin organization, transcription and splicing (Figure 7 and Supplementary Figure 9). Together with the observed differential localisation of mCG and mCA in neuronal nuclei, this data suggests that rather than being directly involved in neuronal specification, mCH could play a role in the dynamic reorganisation of chromatin (Fraser et al., 2015) and the regulation of alternative splicing that occurs during neuronal maturation (Hubbard et al., 2013, Weyn-Vanhentenryck et al., 2018).

## Discussion

The complex dynamics, composition, and patterns of DNA methylation observed in the development and maturation of postnatal neurons is hypothesized to play an important role in modifying gene expression and consolidating both neuronal cell types and their response to activity (Cortes-Mendoza et al., 2013, Day et al., 2013, Feng and Fan, 2009, Graff et al., 2012, Miller and Sweatt, 2007, Stroud et al., 2017). Mouse studies have identified some factors involved in the process, including the DNA methyltransferases catalysing the deposition of mC (Guo et al., 2014, Nguyen et al., 2007), and methylation readers such as MeCP2 that link DNA methylation to gene expression (Chen et al., 2015, Fasolino and Zhou, 2017, Kinde et al., 2016, Lagger et al., 2017, Mellen et al., 2012, Skene et al., 2010, Stroud et al., 2017). However, many basic questions remain unanswered, including the roles and regulation of dynamic methylation events, and the factors that define the targeting of methylation sites. Developing a robust *in vitro* model system to recapitulate the diverse methylation events particular to neurons is important to facilitate the detailed molecular dissection of these processes. In the present study we have extended an established protocol to differentiate cortical neurons from mESCs and shown that these cells acquire *in vivo* levels of non-CG methylation in a similar time frame to *in vivo* brain development. Furthermore, we have shown that the timing of mCH deposition *in vitro* correlates to a transient increase in *Dnmt3a* expression, which also recapitulates that observed *in vivo (Lister et al., 2013)*. If, as these results suggest, the deposition of mCH and *Dnmt3a* expression is indeed hardwired into the developmental process, this has profound implications for studying the equivalent processes in human iPSC-derived neurons. The human brain has a much more extended developmental timeline compared to mouse (Stiles and Jernigan, 2010), and maximal *in vivo* mCH methylation levels are not observed until late adolescence (16+ yr) (Lister et al., 2013). The most advanced *in vitro* human cerebral organoid differentiation protocols currently available can only recapitulate relatively early embryonic developmental stages (reviewed (Benito-Kwiecinski and Lancaster, 2019)). Whether further development of human cerebral organoids through *in vitro* vascularisation (Cakir et al., 2019) or transplantation into the mouse (Mansour et al., 2018) can overcome this developmental obstacle and accelerate human neuron maturation to experimentally tractable time scales is currently unknown. To our knowledge, the mouse data presented here is the first report of mammalian neurons derived *in vitro* from pluripotent cells that harbor mCH at levels similar to those present in neurons *in vivo*, and opens the door to further, targeted investigations of the regulatory pathways and environmental factors involved.

Detailed analysis of DNA methylation levels and genomic distribution in mouse ESC-derived neurons identified several interesting features. Firstly, the levels and distribution of mCG and mCH were regulated independently. Compared to the mouse brain neurons, generalised hypermethylation in the mCG context, both within gene bodies and across the genome, was not observed for mCH, which showed relatively normal gene body methylation levels and slightly lower exon methylation. Regions of higher methylation in CG context were not in general accompanied by higher methylation in CH context and *vice versa*, and regions showing comparably lower methylation in one context did not show the same pattern in the other context. These observations suggest that different regulatory mechanisms are involved in the remodelling of neuron methylation patterns during maturation, depending upon the DNA context and genomic feature targeted. As the mESC-derived neurons did not recapitulate either of these methylation contexts with complete fidelity, it is likely that other factors, such as environment, neuronal connectivity and activity, all act to develop the mature neuronal methylome. Similarly, our comparisons to previously published iN cell data (Luo et al., 2019) suggest that in this model system also, other factors are required to fully develop *in vivo* methylation levels and patterns.

The early post-natal nuclear landscape of neurons is highly dynamic, with changes in gene transcription (Kang et al., 2011), alternative splicing (Furlanis and Scheiffele, 2018, Weyn-Vanhentenryck et al., 2018), DNA methylation (He and Ecker, 2015, Lister et al., 2013, Szulwach et al., 2011), and chromatin remodelling (Gallegos et al., 2018). It is likely that the regulation of all these facets of neuron development are tightly interconnected. It is well-established that gene body mCH levels in neurons is inversely correlated with gene expression (Chen et al., 2015, Chen et al., 2014, Gabel et al., 2015, Lister et al., 2013), and it has been suggested that establishment of early post-natal mCH regulates the transcription of affected genes at later time points (Stroud et al., 2017). Immunologically labelled mCA, both *in vivo* and *in vitro*, shows a strong association with the nuclear periphery, suggesting association of these genomic regions with the nuclear lamina. As the nuclear lamina forms a repressive environment for transcription (Zuleger et al., 2011), association and dissociation of genes with this environment can be a potent regulator of expression. During the neural induction of mESCs, the pro-neural gene *MASH1 is* translocated away from the nuclear lamina, concomitant with upregulation of its expression (Williams et al., 2006). Similarly, hundreds of genes change lamina interactions during differentiation from mESC to neural progenitor cells, and subsequently to astrocytes, and genes affected by altered lamina interactions reflect cell identity and influence the likelihood of a gene being subsequently activated (Peric-Hupkes et al., 2010). These findings suggest that lamina-genome interactions are centrally involved in the control of gene expression programs during lineage commitment and terminal differentiation. The association and dissociation of genes from the nuclear lamina is not restricted to developing or differentiating cells. For example, the *BDNF* gene is translocated away from the nuclear lamina, with a concomitant increase in expression, following stimulation of mature neurons *in vivo*, proving that transcription-associated gene repositioning can occur in adult neurons, as a result of enhanced activity (Walczak et al., 2013). While it is not yet known how mCA associates with the nuclear lamina, or whether this is a direct association, one possibility could involve binding to MeCP2 (Chen et al., 2015, Gabel et al., 2015, Lister et al., 2013, Stroud et al., 2017). MeCP2 is a multifunctional protein, with reported roles in both repression and upregulation of gene expression, as well as involvement in nuclear structure (Chahrour and Zoghbi, 2007, Lagger et al., 2017, Young et al., 2005). In addition to binding various methylated DNA species through its methyl-binding domain, including mCA, mCG, and 5hmC (Chen et al., 2015, Gabel et al., 2015, Guo et al., 2014, Lagger et al., 2017), MeCP2 is able to interact directly with the lamin-B receptor (Guarda et al., 2009), a role that is independent of its function as an epigenetic reader protein. As levels of MeCP2 are very high in neurons, approaching that of H1 linker histone levels (Kishi and Macklis, 2004, Skene et al., 2010), it is tempting to speculate that one role of MeCP2 binding to mCA is to regulate its association with the nuclear lamina. The level of MeCP2 has been shown to increase during mESC-derived neuron differentiation (Yazdani et al., 2012) consistent with a close regulatory association to the increased levels of mCA observed here.

Generation of base resolution DNA methylomes of mESC-derived neurons allowed us to compare pathways for which gene sets showed either the greatest similarity or the greatest difference in methylation levels to *in vivo* neurons. Pathways hypermethylated in the mCH context in ESC-derived neurons relative to *in vivo* neurons included a range of broad developmental pathways, with no particular emphasis on neuron maturation or function. As mCH has been shown to be deposited in the bodies of lowly expressed genes (Stroud et al., 2017), this suggests that these pathways have reduced functionality in post-mitotic ESC-derived neurons. In contrast, pathways with hypomethylation in the mCH context represented the cell cycle and mitosis. These pathways are predicted to be silenced in post-mitotic cells, suggesting that this hypomethylation occurs as a result of the inclusion of the represented genes in tightly packed heterochromatin, although further studies are needed to confirm this. Taken together, these data suggest that despite the different levels and labeling patterns for mCH in the mESC-derived neurons, some fidelity is retained in the ESC-derived cells. Interestingly, gene set enrichment pathways showing the greatest similarity in mCH between *in vivo* and ESC-derived neurons include a range of regulatory pathways involved in transcription, splicing and chromatin organisation, all aspects of neuron development that are significantly modified in early postnatal neurons. Whether mCH is involved in the regulation of alternative splicing or chromatin remodelling in neurons remains to be determined. However, direct links between mCA and alternative splicing have been shown in human ESCs (Tan et al., 2019) and CG methylation contributes to the inclusion or exclusion of alternatively spliced exons in human cell lines (Maunakea et al., 2013). Furthermore, as MeCP2 can also be associated with alternative splicing and nuclear architecture (Yazdani et al., 2012, Young et al., 2005), a potential role for mCH in these processes should be considered.

The generalised CG hypermethylation in mESC-derived neurons challenged the analysis of gene sets enriched for either similarity or difference to *in vivo* neurons, as the highest ranked gene sets for both analyses represented different but closely related aspects of neuron development and function. The mechanism underlying mCG hypermethylation is not yet known. It could reflect either increased methylation, decreased activity of demethylation pathways (Wu and Zhang, 2017), or an accumulation of 5hmC (Lister et al., 2013, Szulwach et al., 2011). or a combination of any or all three. The genome-wide increase in mCG was observed to occur earlier in the neuron differentiation timeline than the maximal increase in mCH, suggesting that these two processes are regulated independently.

Together, this work establishes that *in vitro* differentiation of mouse embryonic cells to neurons is a highly tractable and valuable model system with which to further dissect the roles of DNA methylation and higher order intranuclear architecture on neuron maturation and function.

## Materials and Methods

### Reagents and Antibodies

Primary antibodies are detailed in Supplementary Table 1. Directly-conjugated Alexa488-mouse anti-NeuN was from Millipore (MAB377X). Mouse monoclonal antibodies to 5-methylcytosine-adenosine dinucleoside (mCA) and 5-methylcytosine-guanine dinucleoside (mCG) were raised against KLH-conjugated dinucleosides at the Technology Development Laboratory (Babraham Bioscience Technologies Ltd, Cambridge, UK) (anti-mCA antibody clone 2C8H8A6 currently available from Active Motif, Cat. No 61783/4). Alexa Fluor secondary antibodies were purchased from Life Technologies. ESGRO recombinant mouse Leukaemia Inhibitory Factor (LIF) was obtained from Merck Millipore. The remaining reagents were obtained from ThermoFisher Scientific or Sigma Aldrich unless otherwise specified.

### Mouse Tissue Collection and in vitro neuron differentiation

Adult C57BL/6 mice (8-10 weeks old) were sacrificed and brains dissected and snap frozen on dry ice. Brains were then sectioned on a cryostat at 12 microns, placed on poly-L-lysine slides (VWR) and stored at −80°C until processing. All experi ental procedures were approved by the Animal Welfare, Experimentation and Ethics Committee at the Babraham Institute and were performed under licenses by the Home Office (UK) in accordance with the Animals (Scientific Procedures) Act 1986.

For primary neurons, hippocampal or cortical neurons were cultured from embryonic day 18 C57BL/6 mouse brains as described previously (Lanoue et al., 2017). All procedures were conducted according to protocols and guidelines approved by the University of Queensland Animal Ethics Committee. Isolated E18 neural progenitors were either frozen directly for WGBS (E18 samples) or plated onto 0.1 mg/ml Poly L-Lysine / 8 µg/ml laminin-coated plates in Neurobasal containing 2% B27, 0.5 mM L-glutamine and 1% Pen-Strep and maintained for 14 days. 14DIV neurons were either dissociated with Accutase and frozen as cell pellets for WGBS or washed in PBS and fixed in 2% PFA/PBS for ICC.

### Murine embryonic stem cell culture and in vitro neuron differentiation

Two mESC cells lines, R1 (ATCC SCRC-1011) derived from a 129X1 x 129S1 male blastocyst (Nagy et al., 1993) and G4 derived from a 129S6/SvEvTac x C57BL/6Ncr male blastocyst (George et al., 2007), were maintained on gamma-irradiated mouse embryonic fibroblasts in mESC medium (DMEM) (GIBCO), 15% Hyclone defined FBS (GE Healthcare), 10^3^U/ml ESGRO LIF (Millipore), 1x L-Glutamax, 1x sodium pyruvate, NEAA 1x (Invitrogen), 0.1 mM beta-mercaptoethanol). Cells were fed daily and split based on confluency. As differentiation of the G4 mESC line was slightly more robust this was used routinely, and data shown refers to G4 ESC-derived neurons unless otherwise stated.

Neural differentiation was initiated by dissociating the mESC colonies using Accutase and excess feeder cells were removed from the cell suspensions by panning on gelatin-coated plates. Dissociated cells were counted and transferred to ultra-low attachment cell culture plates at a dilution of 220,000 cells/ml, in differentiation medium (Dulbecco’s odified Eagle Medium (DMEM) (GIBCO), 10% Hyclone defined FBS (GE Healthcare), 1x L-Glutamax, 1x sodium pyruvate, 1x NEAA (GIBCO), and 0.1 mM beta-mercaptoethanol). Neural induction was continued for 8 days, medium was changed every two days, and supplemented with 5 µM retinoic acid for the final 4 days. Cell aggregates were dissociated using Accutase, single cell selected through a 40 µm cell strainer, and plated onto 0.1 mg/ml poly-ornithine / 4 µg/ml laminin-coated plates in N2 medium (DMEM/F12, 1x Glutamax 1x N2 supplement, 20 µg/ml insulin, 50 µg/ml BSA) at a density of 50,000-100, 000 cells/cm^2^. After 2 days, cells were changed into N2B27 (Neurobasal, 1x N2 supplement, 1x B27 supplement, 1x Glutamax). N2B27 was changed every 4 days for 12 days, following which 50% media changes were performed for the remaining culture time.

High potassium-mediated depolarisation was performed as previously described (Martin et al., 2014). Briefly, cells were washed with low K+ buffer (15 mM HEPES, 145 mM NaCl, 5.6 mM KCl, 2.2 mM CaCl_2_, 0.5 mM MgCl_2_, 5.6 mM D-glucose, 0.5 mM ascorbic acid, 0.1% BSA, pH 7.4), transferred to high K+ buffer for 5 min (formulation as the low K^+^ buffer but using 95 mM NaCl and 56 mM KCl), washed again in low K+ buffer and either harvested directly for RNA, immediately fixed for ICC, or chased in growth medium for the appropriate times prior to fixation or RNA extraction.

### Human in vitro neuron differentiation

Human neurons were generated following two different approaches: either as adherent monolayer cultures via a neural stem cell intermediate population (Reinhardt et al., 2013) or as cerebral organoids (Lancaster and Knoblich, 2014). For adherent cultures, PSCs were disaggregated mechanically and cultivated as embryoid bodies in DMEM/F12 and Neurobasal 1;1 mix, 1% Glutamax (all Life Technologies) with 1:200 N2 Supplement (R&D Systems), 1:100 B27 supplement without retinoic acid (Miltenyi Biotec) supplemented with 10 µM SB-431542, 1 µM dorsomorphin (both Selleckchem), 3 µM CHIR 99021 (Cayman Chemical Company) and 0.5 µM PMA (Sigma) for 3 days on petri dishes. On day 4, SB-431542 and dorsomorphin were removed and 150 µM ascorbic acid (Cayman Chemical Company) was added to the media. On day 6, cells were plated onto Matrigel-(Corning) coated TC dishes to allow attachment of neuroepithelial cell types. Over several passages, neural stem cells were enriched resulting in pure populations. These neural stem cells could be differentiated by removing the small molecules from the media and switching to B27 supplement with retinoic acid, resulting in a mixed population of beta3-Tubulin positive neurons and GFAP-positive glia within 4 weeks. For cerebral organoids, PSCs were dissociated into single cells and plated in a ultra-low-attachment 96 well plate using 9,000 cells/ well in E8 media (Life Technologies), with 50 µM Y-27632 (Selleckchem). After 5-6 days, embryoid bodies were transferred to ultra-low-attachment 6-well TC plates in neural induction media (DMEM/F12, 1% Glutamax, 1% non-essential amino acids and 10 µg/ml heparin (Sigma). After another 4-5 days, organoids were embedded into Cultrex (R&D Systems) matrix and cultivated under continuous agitation on an orbital shaker in cerebral organoid media (DMEM/F12 and Neurobasal 1;1 mix, 1% Glutamax, 0.5% non-essential amino acids, 1:200 N2 Supplement, 1:100 B27 supplement (Miltenyi Biotec) and 1:40,000 Insulin (Sigma), with media changes every 3 days until a maximum age of 9 months.

### RT-qPCR

Total RNA was extracted using the Macherey-Nagel NucleoSpin RNA kit, including on column digestion of DNA with RNase-free ase according to anufacturer’s specifications. Concentration and 260/280 ratios were quantified using a NanoDrop 1000 spectrophotometer before cDNA synthesis using the iScript cDNA synthesis kit (Bio-Rad). Primers were designed to span exon-exon boundaries wherever possible (Supplementary Table 2). When this was not possible, samples were excluded if genomic DNA contamination was more than 10-fold over the cDNA concentration. Quantitative PCR (qPCR) reactions used SsoFast Evagreen (Bio-Rad) with c A te plate according to anufacturer’s instructions, using a C1000 Thermocycler (Bio-Rad) and CFX software. Results were analysed as described previously (Livak and Schmittgen, 2001).

### Immunocytochemistry

Protein ICC was performed as described previously (Martin et al., 2013) on cells fixed in 4% paraformaldehyde or 2% paraformaldehyde (Electron Microscopy Sciences). For double and triple immunocytochemistry using antibodies to methylated DNA, an alternative sequential labelling method was used. Briefly, cells were fixed in 2% PFA/PBS for 10-30 min, permeabilised for 1 h in 0.5% TX-100, then depleted of residual methylated RNA using RNase A at 10 µg/ml for 30 min at 37°C in PBS. Cells were subsequently blocked in PBS/0.2% BSA/0.2% cold-water fish skin gelatin for 10 min, then labelled with protein-targeting antibodies in PBS/0.2% BSA/0.2% cold-water fish skin gelatin overnight at 4°C. Cells were then washed with PBS, labelled using corresponding secondary antibodies in PBS/0.2% BSA/0.2% cold-water fish skin gelatin for 30 min, washed in PBS, then re-fixed in 2% PFA/PBS for 15 min. DNA epitopes were retrieved using 4N HCl/0.1% TX-100 for 10 min at room temp, cells washed in PBS/0.05% Tween-20, and blocked in PBS/0.05% Tween-20/1% BSA (BS) for 1 h. Antibodies to methylated DNA were applied in PBS/0.05% Tween-20/1% BSA overnight at 4°C. Cells were subsequently washed in PBS/0.05% Tween-20/1% BSA and appropriate secondary antibodies applied in the same buffer containing 0.1 µg/ml DAPI for 30 min-1 h. Finally, cells were washed in PBS/0.05% Tween-20/1% BSA, rinsed in PBS and either imaged directly (high throughput imaging), or mounted in Mowiol (Confocal).

### Immunohistochemistry

Mouse brain sections were fixed in 2% PFA for 30 min, then permeabilised in PBS/0.5% Triton X-100 for 1h, blocked in BS for 1h and incubated overnight at 4°C with a NeuN antibody (1:2500) in BS. They were subsequently incubated for 1h with a secondary fluorescently labelled antibody in BS (1:1000). After post-fixation with 2% PFA for 10 min, to obtain access to DNA methyl groups, cell nuclei were mildly depurinised with 4N HCl treatment for 15 min and incubated with antibody-enriched culture medium against mCG or mCA in BS overnight at 4°C. After incubation for 1h with a secondary fluorescently labelled antibody in BS (1:1000) the tissue was stained with 1:100 YOYO1 (Molecular Probes, Life Technologies) for 15 min and mounted with Vectashield Antifade Mounting Medium (H-1000, Vector Laboratories). Washing with PBS/0.05% Tween-20 for 1-2h was performed after each step, reagents were dissolved in PBS/0.05% Tween-20 and steps performed at room temperature, unless otherwise stated.

### Microscopy

Phase contrast microscopy was performed on live cells using an Olympus IX51. Confocal microscopy was performed on fixed cells using either a Zeiss 710 confocal microscope and a 40x water immersion objective or a Leica SP8 confocal microscope using a 60x oil immersion objective. High throughput imaging was performed using a Perkin-Elmer Operetta equipped with a 20x Air objective. Brain section imaging was performed with Nikon A1-R confocal microscope and a 60x 1.4 NA oil immersion objective.

Image analysis was performed by manual masking of nuclei and measuring fluorescence intensity/nucleus using Image J (Figure 1J, L) or by high-throughput Operetta image acquisition using Harmony (Figure 1D, F, H; Supplementary Figure 1E; Supplementary Figure 2B), prior to exporting the images, generating nuclear masks and analysing fluorescence intensity using Cell Profiler (Carpenter et al., 2006). All image analysis data was collated using Excel 2016, and graphs prepared using Excel for Office 365 or Graphpad Prism 8. All images were processed using Adobe Photoshop 2020 and figures compiled with Adobe Illustrator 2020.

### Transmission electron microscopy

ESC-derived neurons were incubated with 10 µg/ml CTB-HRP in either high K^+^ or low K^+^ buffer for 5 min, washed in PBS and fixed in 2.5% glutaraldehyde (Electron Microscopy Sciences) for 24 h. Following fixation, cells were processed for 3, 39-diaminobenzidine (DAB) cytochemistry using standard protocols. Fixed cells were contrasted with 1% osmium tetroxide and 4% uranyl acetate prior to dehydration and embedding in LX-112 resin (Martin et al., 2013). Sections (∼50 nm) were cut using an ultramicrotome (UC64; Leica). To determine CTB-HRP endocytosis, presynaptic regions were visualized at 60,000x using a transmission electron microscope (model 1011; JEOL) equipped with a Morada cooled CCD camera and the iTEM AnalySIS software.

### Whole genome bisulfite sequencing

DNA methylation analysis by WGBS was performed using approximately 100,000 cells. Genomic DNA was isolated with the DNeasy Blood and Tissue Kit (Qiagen) with some modifications: samples were incubated for 4 h at 56°C with an additional RNAse A incubation for 30 min at 37°C. 500 ng of genomic DNA spiked with 4% (w/w) unmethylated lambda phage DNA (Thermo Fisher Scientific) was sheared to a mean length of 200 bp using the Covaris S220 focused-ultrasonicator. Libraries for WGBS were prepared as follows: DNA fragments were end-repaired using the End-It kit (Epicentre), A-tailed with Klenow exo-(NEB) and ligated to methylated Illumina TruSeq adapters (BIOO Scientific) with DNA Ligase (NEB), followed by bisulfite conversion using EZ DNA-methylation Gold kit (Zymo Research). Library fragments were then subjected to 7 cycles of PCR amplification with KAPA HiFi Uracil+ DNA polymerase (KAPA Biosystems). Single-end 100 bp sequencing was performed on a HiSeq 1500 or a MiSeq (Illumina).

### DNA methylation analysis

Reads were trimmed for quality and adapter sequences removed. Following pre-processing, reads were aligned to mm10 or hg19 references with Bowtie and a pipeline described previously (Langmead et al., 2009, Lister et al., 2009), resulting in a table summarising methylated and unmethylated read counts for each covered cytosine position in the genome. Bisulfite non-conversion frequency was calculated as the percentage of cytosine base calls at reference cytosine positions in the unmethylated lambda control genome. This was performed individually for each context (CA, CC, CG, CT). Methylation for particular genomic contexts and average methylation of gene bodies were calculated by intersecting whole genome data with feature bed files from the UCSC table browser using Bedtools (Quinlan, 2014). For correlation, heatmap generation and clustering of methylation data, Deeptools was used (Ramirez et al., 2016), along with R using packages pals, gplots, ggplot2, viridis and RColorBrewer. When calculating average methylation level in a region (weighted methylation level), the number of C basecalls divided by total sequence coverage at reference C positions was used to calculate the methylation level for each context (CG, CH, CA, CC, CT). Sample-specific and context-specific bisulfite non-conversion rates, calculated from the unmethylated lambda phage DNA control, were subtracted for each methylation context. Gene set enrichment analysis was performed using the fgsea package for R (Subramanian et al., 2005) with reactome (Fabregat et al., 2018) and gene ontology annotations (Ashburner et al., 2000, The Gene Ontology, 2019). Pathways were sorted by NES score (enrichment score normalized to mean enrichment of random samples of the same size) and only pathways with p-value < 0.05 were considered in subsequent analyses.

For similarity analysis, every gene was given a score as follows: First, the difference in weighted DNA methylation between *in vivo* and *in vitro* neurons (y) was calculated for each gene and scaled to a value between 0 and 1 (using the formula x = |y-1|), so that genes that were more similar in methylation state between both samples would have a value (x) closer to 1. Then, the average of weighted DNA methylation per gene in neuronal samples was compared against the average weighted DNA methylation of glial and fetal samples, resulting again in a score between 0 and 1, with genes showing greater differences having a value closer to 1. Both scores for similarity between neurons and dissimilarity to non-neuronal samples were added in order to give both aspects the same weight, and scaled to values between 0 and 1, with values closer to 1 representing genes being more similar in methylation state between both neuronal samples but different compared to glia and fetal frontal cortex. This similarity score was then used to rank all genes for GSEA using the fgsea package as described above.

### Fluorescence-activated nuclear sorting

Intact nuclei were isolated from cell pellets as described previously (Li et al., 2014, Okada et al., 2011). Briefly, cells were Dounce-homogenised on ice in chilled nuclear extraction buffer (10 mM Tris-HCl, pH 8, 0.32 M sucrose, 5 mM CaCl_2_, 3 mM Mg(Ac)_2_, 0.1 mM EDTA, 1 mM DTT, 1x protease inhibitor cocktail (Merck), 0.3% Triton X-100). Nuclear lysates were filtered (40 µm), centrifuged for 7 min at 3000 rpm at 4°C and resuspended in PBS. Nuclei were blocked with 10% normal goat serum and labelled for 60 min on ice with either directly conjugated mouse anti-NeuN-Alexa488, or pre-conjugated rabbit anti-Nanog/goat anti-rabbit Alexa488 or rabbit anti-Pax6/goat anti-rabbit Alexa488 complexes. Samples of each nuclear fraction were retained for secondary only antibody controls. 7-AAD (20 µg/ml) was added all samples 15 min prior to sorting. A BD Influx cell sorter was used to sort nuclei. Prior to sorting, the secondary only control was used to gate events to isolate nuclei from cell debris. From the selected nuclear populations, nuclei were separated into distinct NeuN+ve/7-AAD+ve and NeuN-ve/7-AAD+ve populations, Nanog+ve/7-AAD+ve and Pax6+ve/7-AAD+ve populations, depending upon the cell type.

**Table 1.** Antibodies.

| Target | Host | Clone | Supplier | Catalog # |
| --- | --- | --- | --- | --- |
| TUBB3 (Beta III Tubulin) | Rabbit |  | Sigma-Aldrich | T2200 |
| TUBB3 (Beta III Tubulin) | Chicken |  | Merck-Millipore | AB9354 |
| NeuN (ICC) | Rabbit | D4G4O | Cell Signaling Technology | 24307 |
| NeuN (IHC) | Rabbit |  | Abcam | ab128886 |
| NeuN-Alexa488 | Mouse | A60 | Merck-Millipore | MAB377X |
| c-Fos | Guinea pig |  | Synaptic Systems | 226 004 |
| Synapsin 1 | Rabbit |  | Merck-Millipore | AB154 |
| Pax6 | Rabbit | Poly19013 | BioLegend | 19013 |
| 5-mC | Rabbit | RM231 | Abcam | ab214727 |
| 5-hmC | Rabbit |  | Active Motif | 39769 |
| mCA | Mouse | 2C8H8A6 | Available through Active Motif | 61783/4 |
| mCG | Mouse | 3A7 | Reik Laboratory, Babraham Institute | - |
| GFAP | Rabbit |  | Dako | Z0344 |
| Nanog | Rabbit | D2A3 | Cell Signaling Technology | 8822 |

**Table 2.**
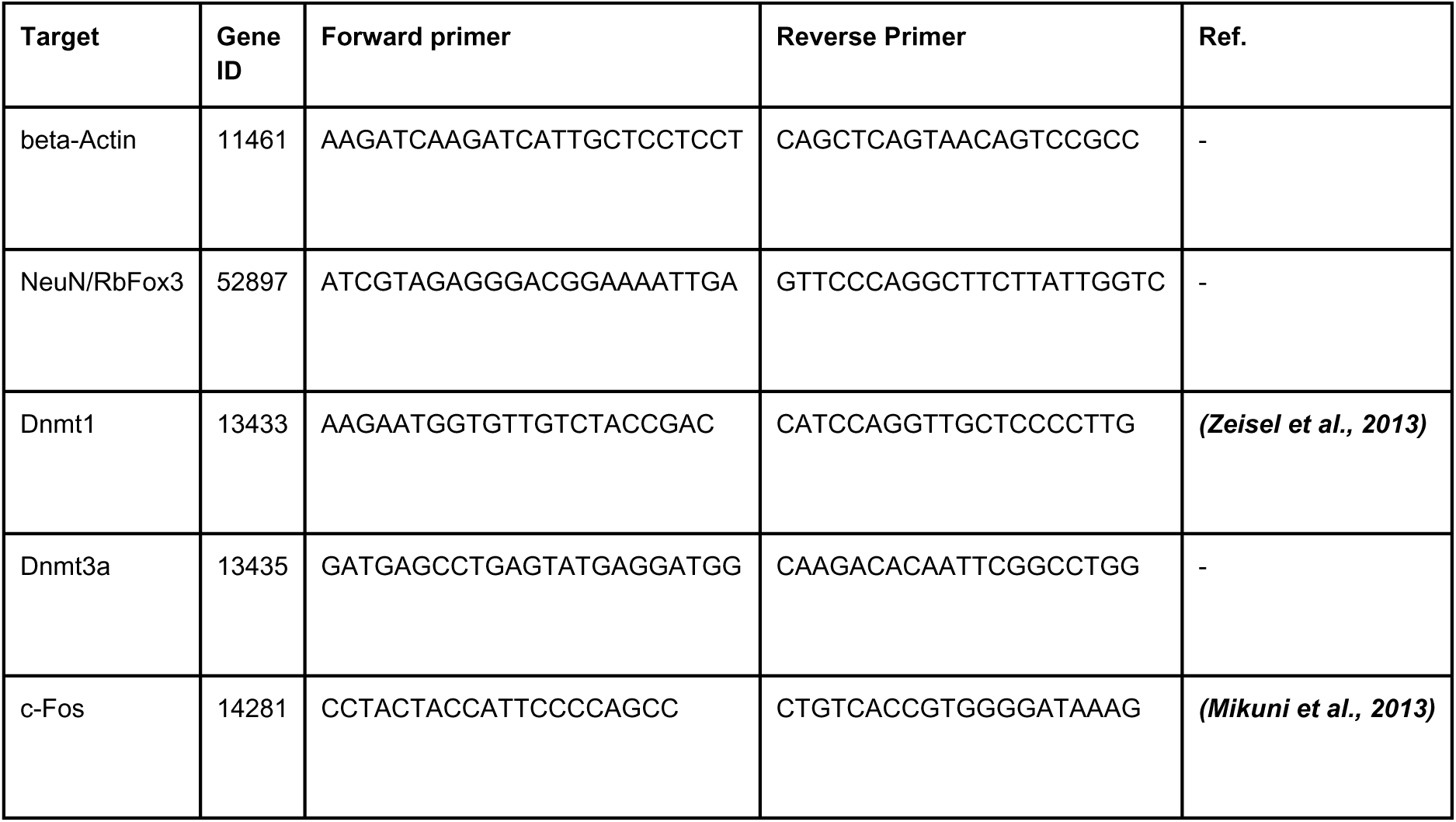
RT-qPCR primers.

### Statistical Analysis

All data was analysed using an unpaired, two-tailed Student’s t-test, unless stated otherwise.

## Acknowledgements

Funding: Monoclonal antibody generation and N.O. were funded by the Babraham Institute Knowledge Exchange and Commercialisation Fund, W.R. is funded by the BBSRC. E.W. was funded by NHMRC APP1130168 and NHMRC APP1090116 project grants. R.L. was supported by a Sylvia and Charles Viertel Senior Medical Research Fellowship and Howard Hughes Medical Institute International Research Scholarship, and research activities by NHMRC GNT1130168 and GNT1090116 Project Grants.

Methylated oligonucleotide sequences were kindly provided by Dr Akanksha Singh, Active Motif (Carlsbad, CA). KLH-conjugated dinucleotides for monoclonal antibody generation were kindly provided by Active Motif Inc. The R1 and G4 mESC cell lines were provided by Prof Peter Koopman (IMB, UQ) and Dr Josephine Bowles (SBMS, UQ). Electron microscopy was performed at the Australian Microscopy and Microanalysis Facility at the Centre for Microscopy and Microanalysis at the University of Queensland. Confocal microscopy (apart from Nikon A1-R) was performed at the Queensland node of the Australian National Fabrication Facility, a company established under the National Collaborative Research Infrastructure Strategy to provide nano- and micro-fa rication facilities for Australia’s researchers. Fluorescent-activated nuclear sorting was performed by the Queensland Brain Institute Flow Cytometry Facility. We thank Lidia Madrid and Zukrofi Muzar for expert technical assistance and Gavin Kelsey (Babraham Institute, UK) for providing reagents. We thank Saskia Freytag for her assistance with R scripting. We thank Marga Behrens for insightful discussions on DNA methylation dynamics during brain development.

## Data availability

WGBS data is available in GEO under the accession number GSE137098.

---

**Supplementary Figure 1.**
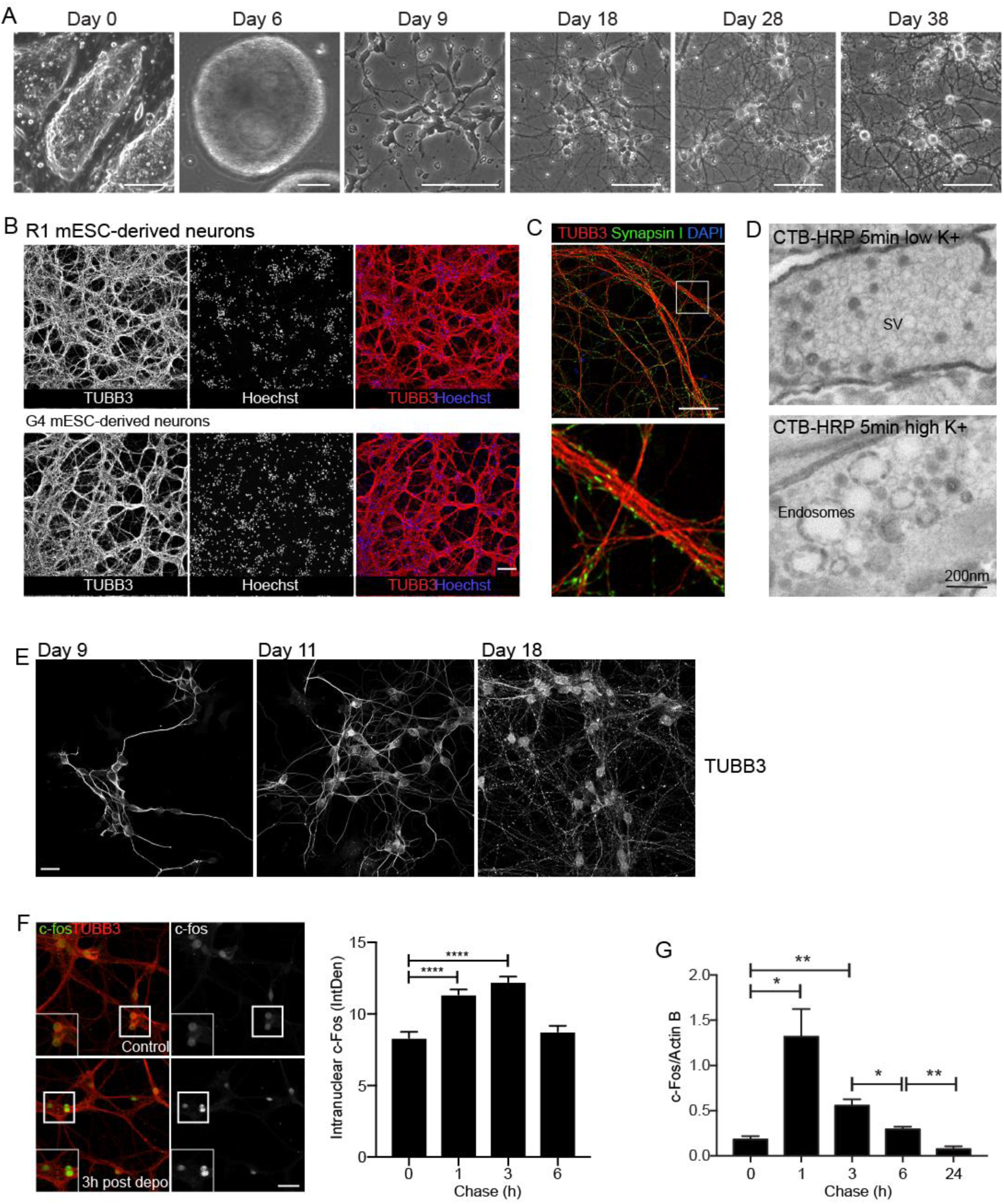
Characterisation of mESC-derived neurons. **(A)** Phase contrast images of mESCs growing on a layer of feeder MEFs, cell aggregates (day 6), neural progenitors (day 9), and neurons (day 18-38). Scale bar =100 µm, except for cell aggregates = 200 µm. **(B)** Neurons derived from either G4 mESCs or R1 mESCs were fixed and labelled for the pan-neuronal marker beta3-tubulin (TUBB3). Both cell lines generated complex neurite networks within 38 days. Scale bar = 100 µm. **(C)** Neurons derived from G4 mESCs were fixed and labelled for TUBB3 and the pre-synaptic protein synapsin 1. Punctate labelling for Syn1 along neurites demonstrates the presence of nascent synapses. Scale bar = 50 µm. **(D)** The response of synapses to depolarisation was shown by transmission electron microscopy. R1 mESC-derived neurons were incubated for 5 min in the presence of CTB-HRP in either low K^+^ or high K^+^ buffer. Low levels of tracer endocytosis into synaptic vesicles in low K^+^ was superseded by high levels of bulk endocytosis in depolarised cells, suggesting a strong rapid burst of neuroexocytosis and compensatory endocytosis (Cousin, 2009). **(E)** G4 mESC-derived neurons were fixed and labelled for TUBB3 at different time points during differentiation. Scale bar = 20µm. **(F)** G4 mESC-derived neurons were depolarised for 5 min with high K^+^, then chased in growth medium for between 1h and 6 h. Cells were fixed and labelled for c-Fos and TUBB3. The intranuclear intensity (IntDen) of c-Fos was determined during the chase. Results = mean ± SEM, for one representative differentiation. Similar labelling profiles were observed in two separate differentiations. Depolarisation resulted in a transient increase in intranuclear c-Fos labelling in the post-depolarisation period. **** p <0.0001, Student’s t-test. Scale bar = 50 µm. **(G)** c-Fos mRNA abundance was determined by RT-qPCR in control G4 mESC-derived neurons and during a 24 h chase following a 5 min transient depolarisation by high K^+^. Results shown are mean ± SEM relative to beta-actin levels. N=3 independent experiments. * p <0.05, ** p <0.01, Student’s t-test.

**Supplementary Figure 2.**
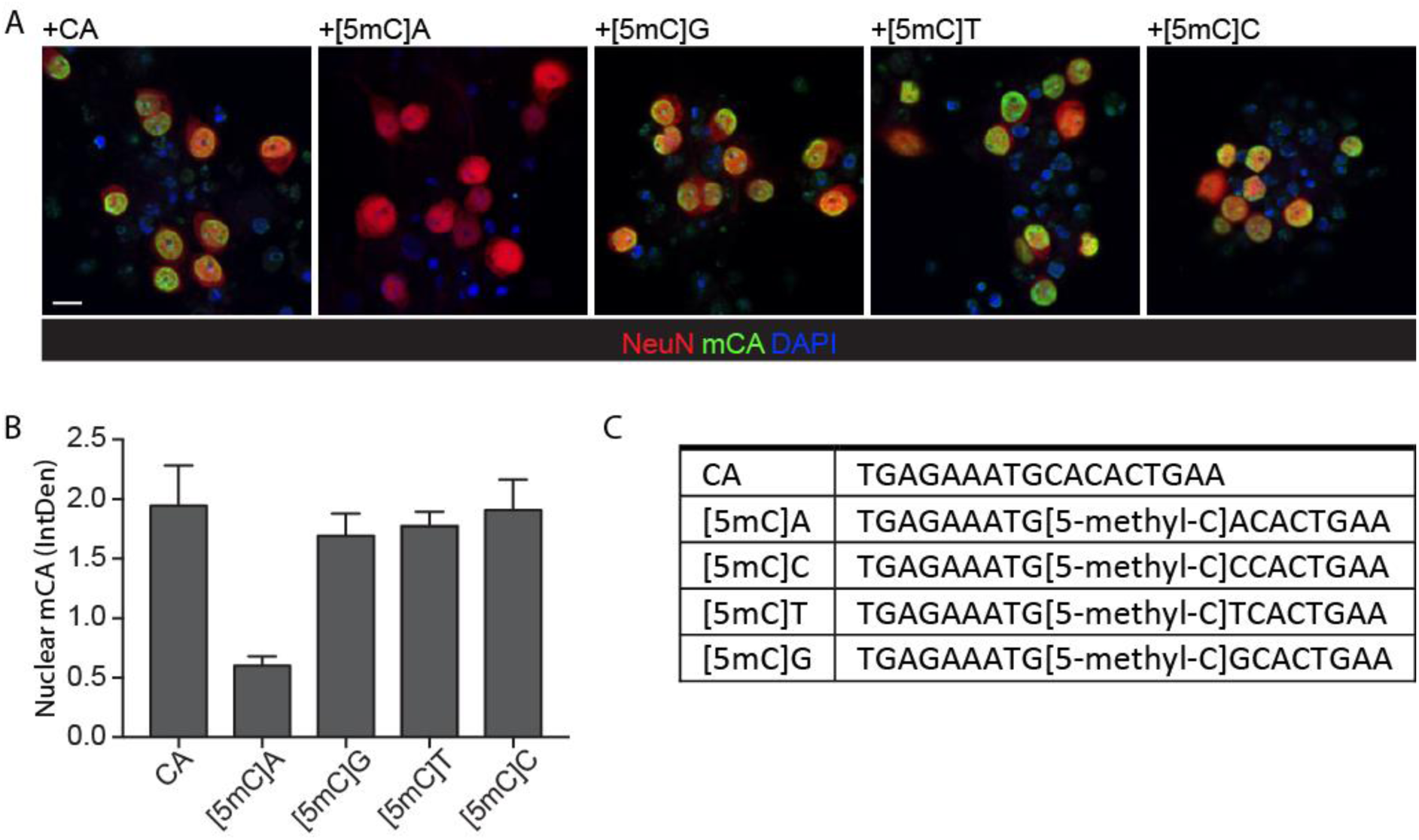
Specificity of the anti-mCA antibody determined by ICC. **(A)** mESC-derived neurons were fixed and immunolabelled for NeuN and mCA ± 2.5 µM competitive methylated oligonucleotides: [5mC]A, [5mC]G, [5mC]T and [5mC]C, or the non-methylated CA oligonucleotide. Scale bar = 10µm. **(B)** The Integrated Density (IntDen) of the nuclear mCA labelling in NeuN-masked nuclei was determined. **(C)** Sequence of the methylated oligonucleotides.

**Supplementary Figure 3.**
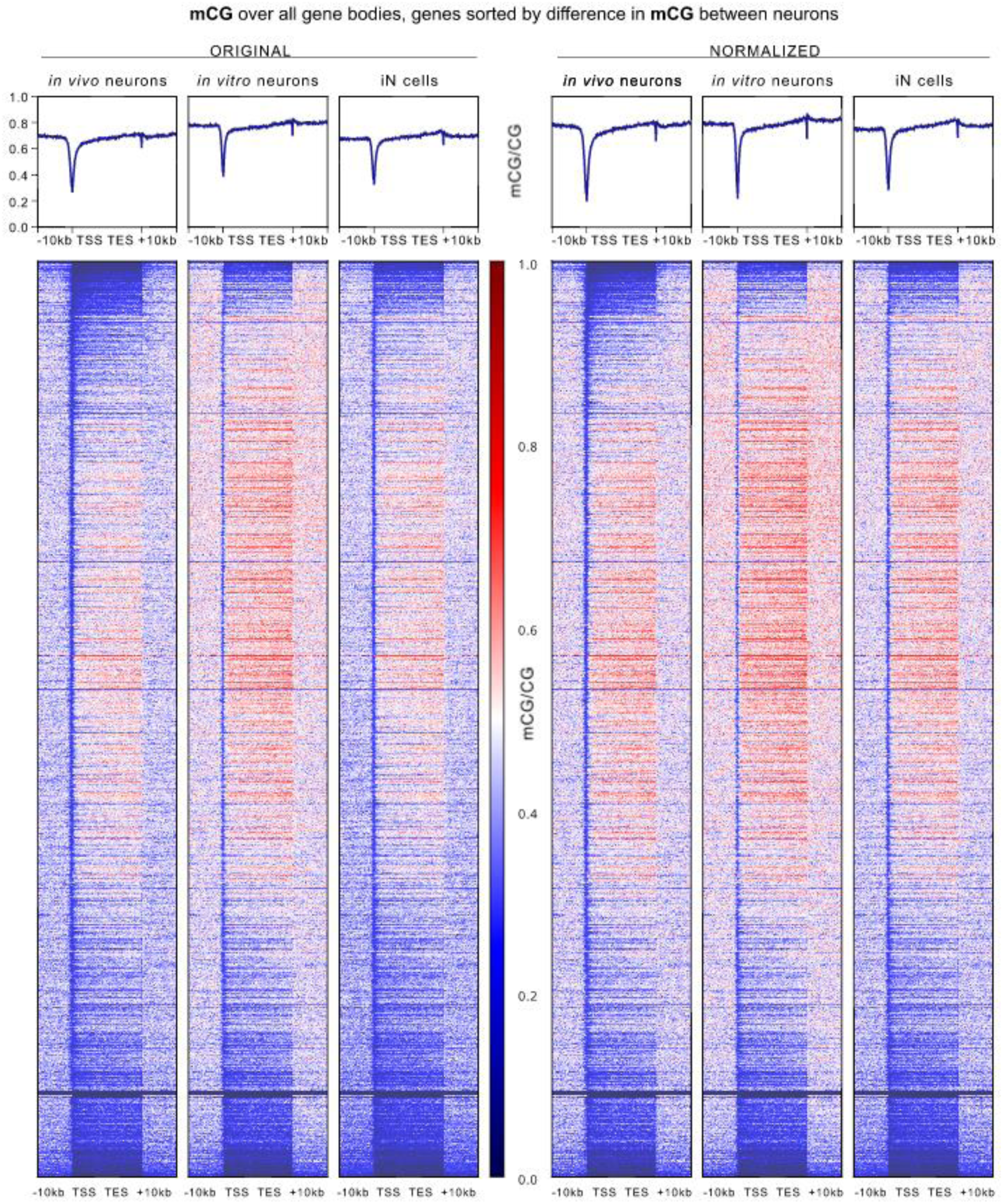
DNA methylation in CG context in gene bodies in ESC-derived and iN cells compared to in vivo neurons. Genes in the same order based on CG methylation difference as in Fig 5A but showing CG methylation for gene bodies and flanking 10 kb for 7-week adult mouse prefrontal cortex neurons (in vivo neurons), mESC-derived neurons (in vitro neurons) and trans-differentiated neurons (iN cells) as reported by (Luo et al., 2019). Left side = original data, right side = normalized to average global CG methylation levels.

**Supplementary Figure 4.**
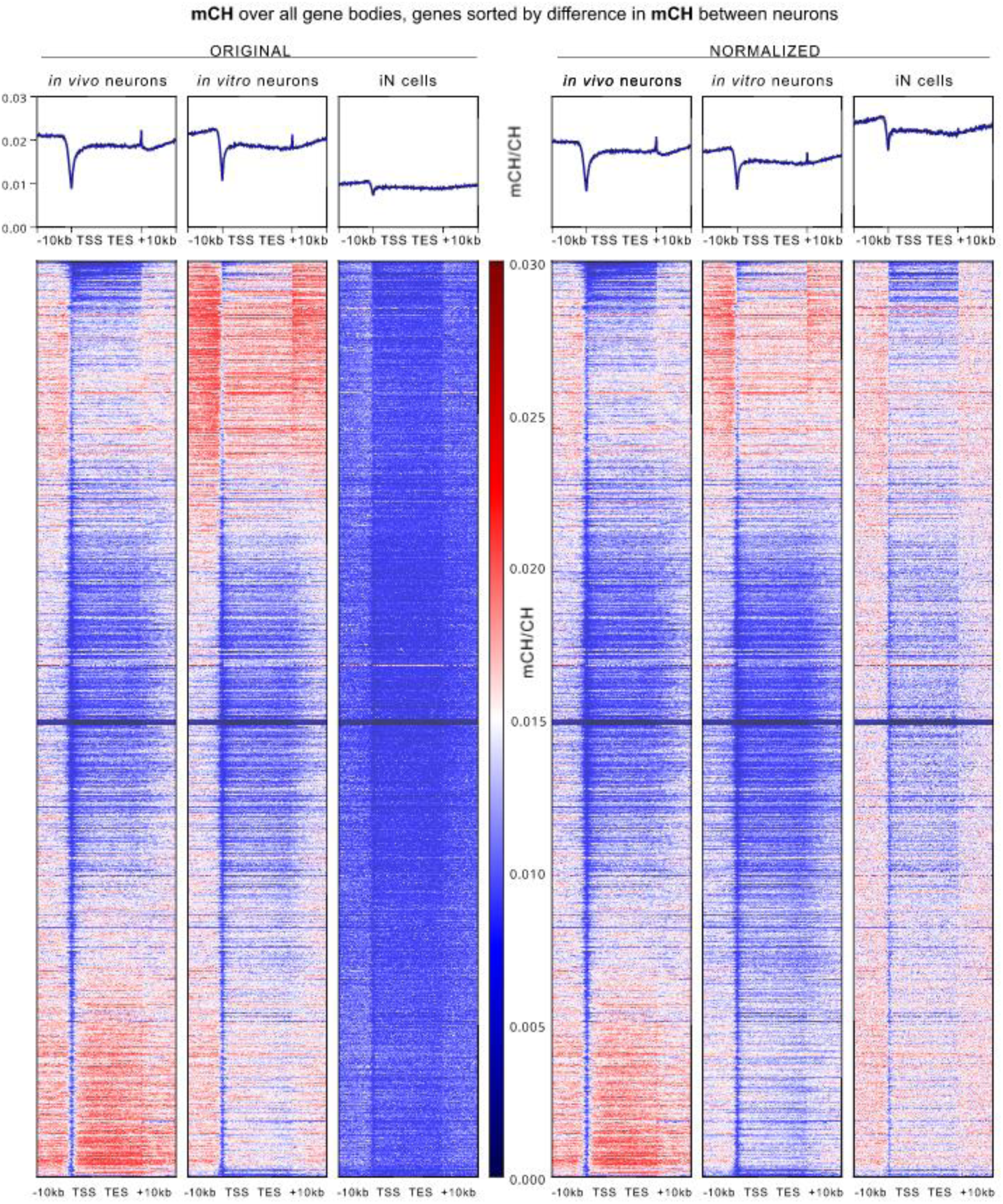
DNA methylation in CH context in gene bodies in ESC-derived and iN cells compared to in vivo neurons. Genes in the same order based on CH methylation difference as in Fig 6A but showing CH methylation for gene bodies and flanking 10 kb for 7-week adult mouse prefrontal cortex neurons (in vivo neurons), d38 mESC-derived neurons (in vitro neurons) and trans-differentiated neurons made from fibroblasts (iN cells) as reported by (Luo et al., 2019). Left side = original data, right side = normalized to average global CH methylation levels.

**Supplementary Figure 5.**
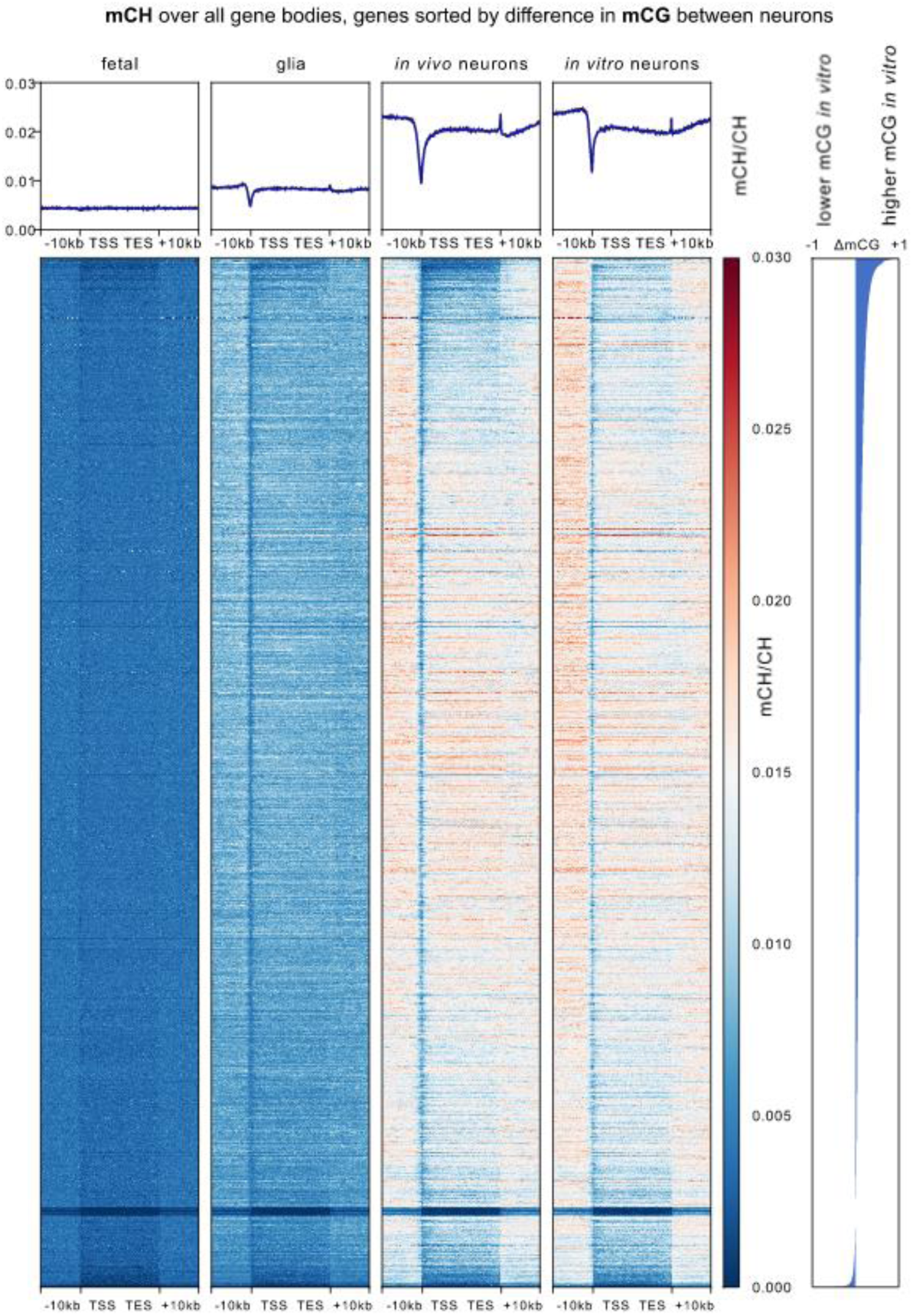
DNA methylation in CH context in gene bodies sorted for differences in mCG between neuronal samples. Genes in the same order based on CG methylation difference as in Fig 5A but showing CH methylation for gene bodies and flanking 10 kb for fetal mouse frontal cortex (fetal), NeuN-negative cells from 7-week adult mouse prefrontal cortex (glia), 7-week adult mouse prefrontal cortex neurons (in vivo neurons), and d38 mESC-derived neurons (in vitro neurons). Difference in mCG between both neuronal samples used for gene order is shown on the right

**Supplementary Figure 6.**
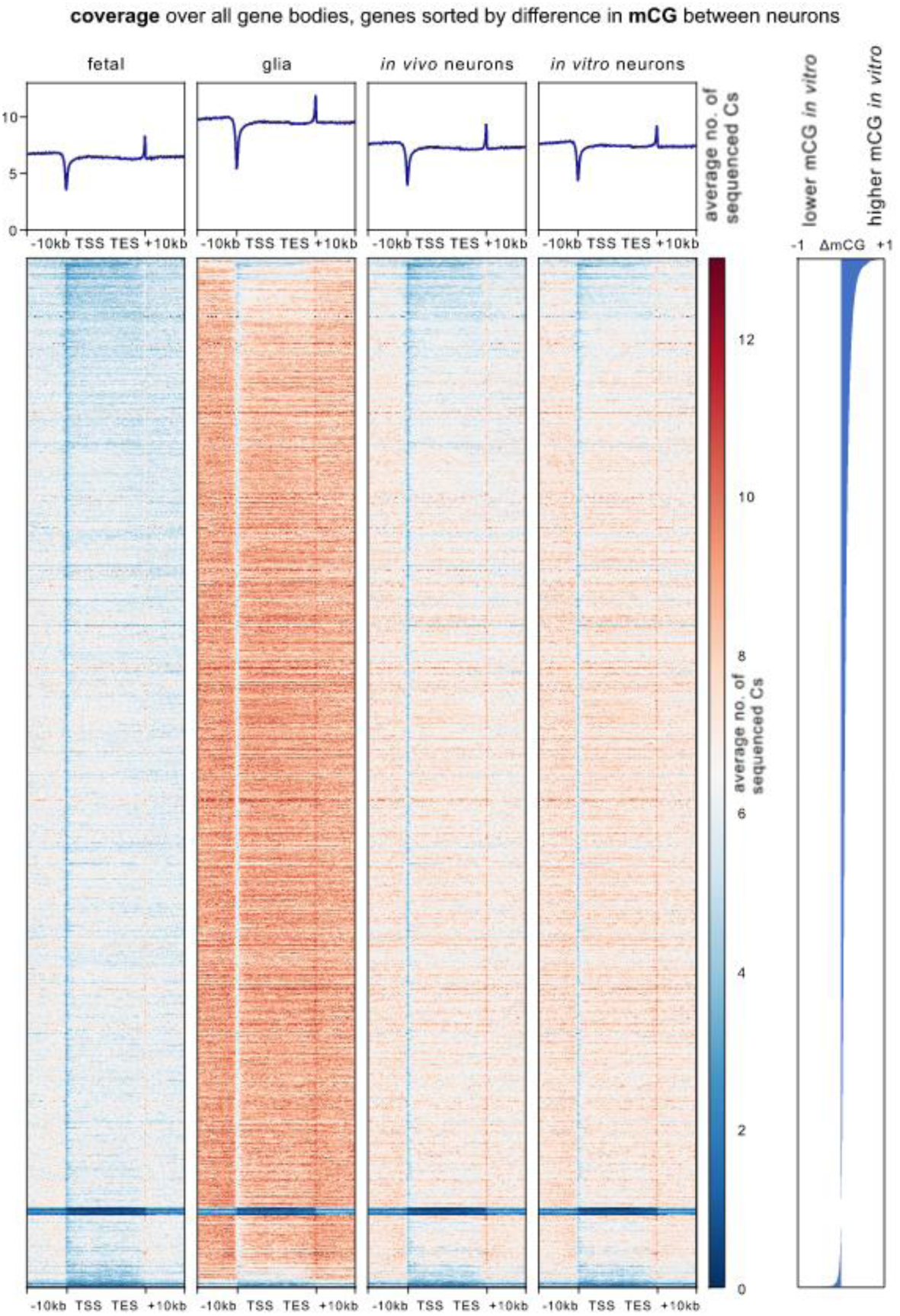
Cytosine coverage in gene bodies sorted for differences in mCG between neuronal samples. Genes are in the same order based on CG methylation difference as in Fig 5A but showing average number of covered cytosines per bin for gene bodies and flanking 10 kb for fetal. mouse frontal cortex (fetal), NeuN-negative cells from 7-week adult mouse prefrontal cortex (glia), 7-week adult mouse prefrontal cortex neurons (in vivo neurons), and d38 mESC-derived neurons (in vitro neurons). Difference in mCG between both neuronal samples used for gene order is shown on the right.

**Supplementary Figure 7.**
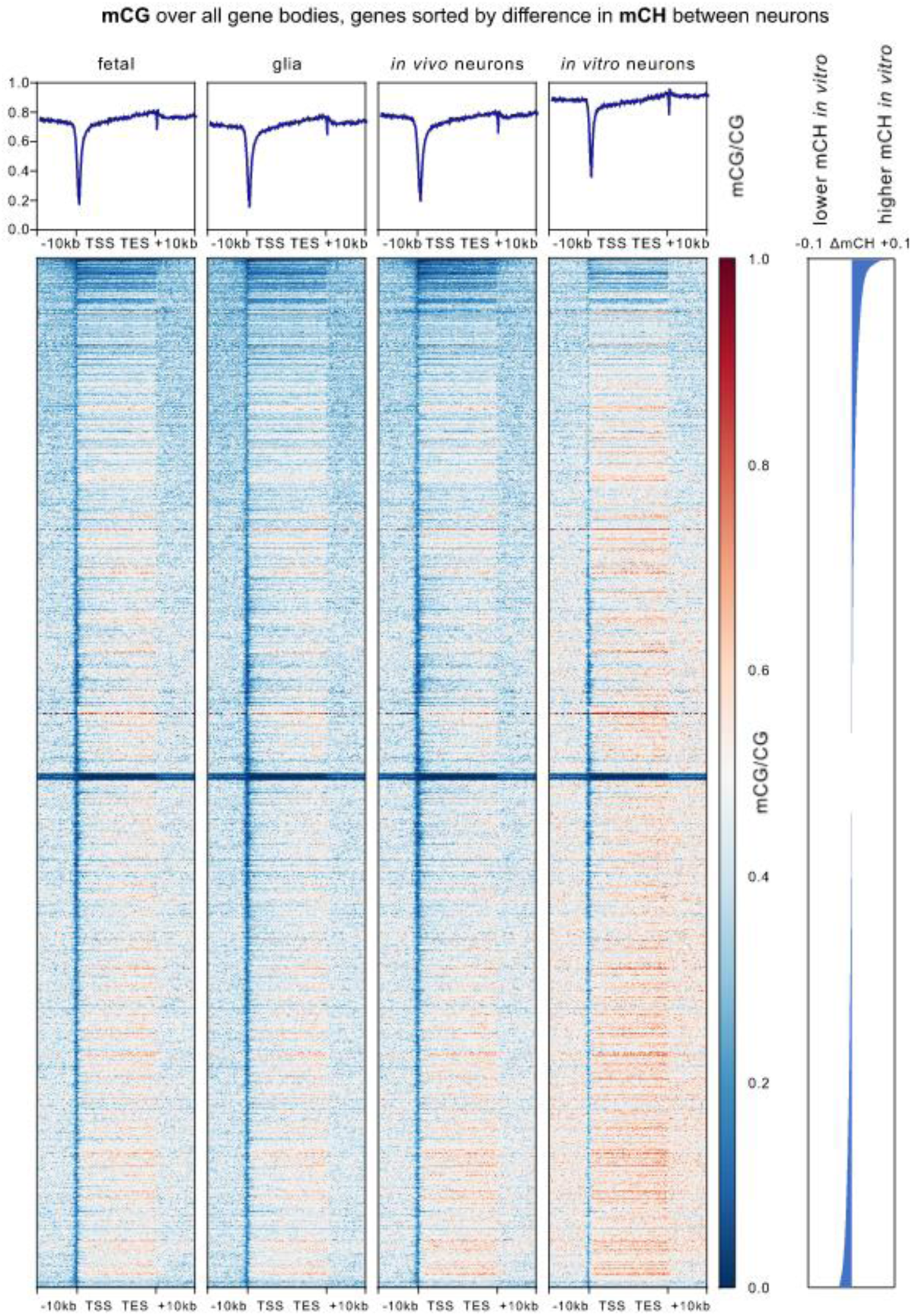
DNA methylation in CG context in gene bodies sorted for differences in mCH between neuronal samples. Genes are in the same order based on CH methylation difference as in Fig 5B but showing CG methylation for gene bodies and flanking 10 kb for fetal mouse frontal cortex (fetal), NeuN-negative cells from 7-week adult mouse prefrontal cortex (glia), 7-week adult mouse prefrontal cortex neurons (in vivo neurons), and d38 mESC-derived neurons (in vitro neurons). Difference in mCH between both neuronal samples used for gene order is shown on the right.

**Supplementary Figure 8.**
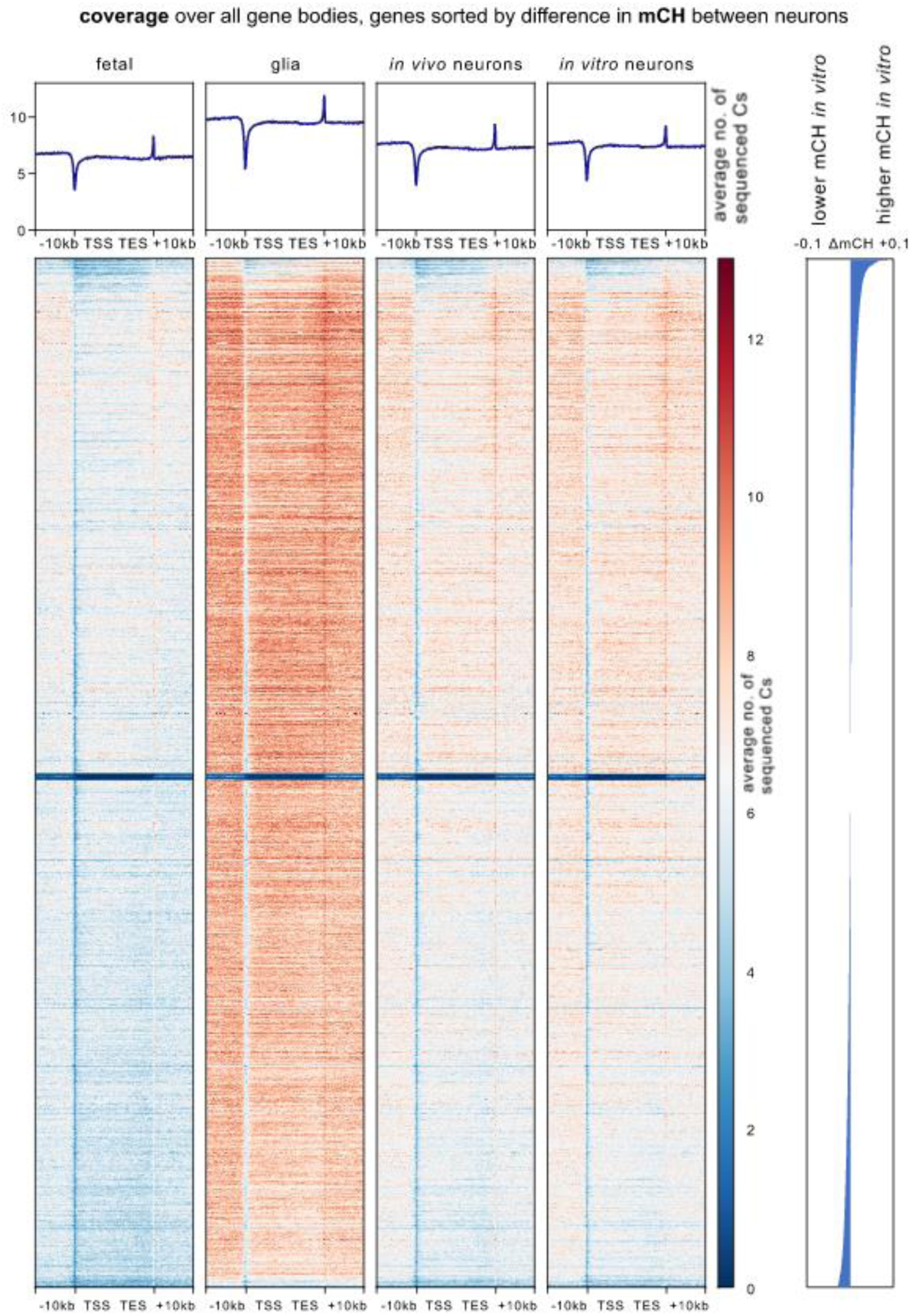
Cytosine coverage in gene bodies sorted for differences in mCH between neuronal samples. Genes are in the same order based on CH methylation difference as in Fig 5B but showing average number of covered cytosines per bin for gene bodies and flanking 10 kb for fetal mouse frontal cortex (fetal), NeuN-negative cells from 7-week adult mouse prefrontal cortex (glia), 7-week adult mouse prefrontal cortex neurons (in vivo neurons), and d38 mESC-derived neurons (in vitro neurons). Difference in mCH between both neuronal samples used for gene order is shown on the right.

**Supplementary Figure 9.**
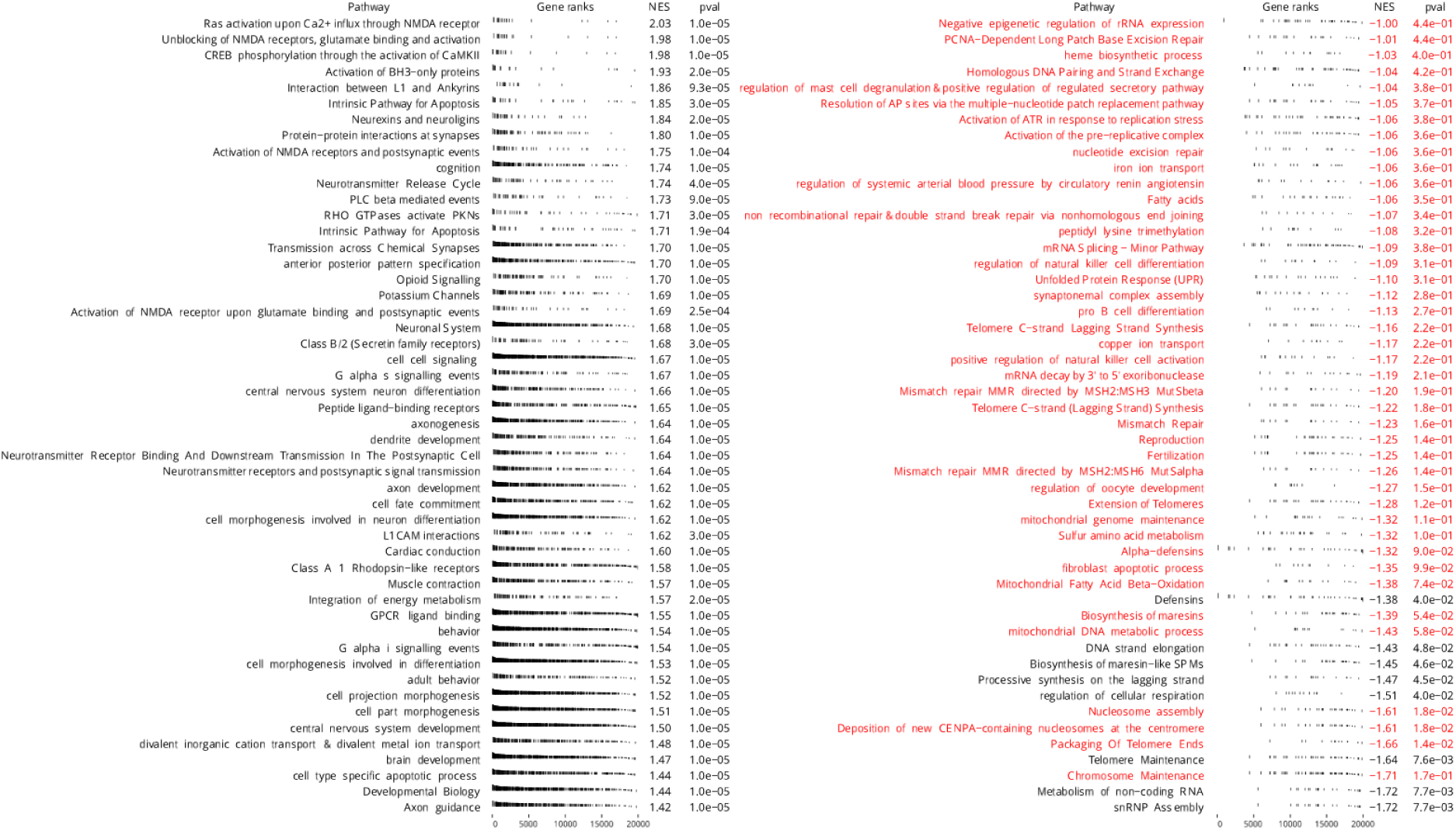
Top 50 enriched pathways for genes differentially methylated in CG context between in vitro neurons and in vivo neurons. Pathways were ranked by enrichment score (NES), whereas positive NES indicates pathways enriched in genes hypermethylated for CG in in vitro neurons, while negative NES indicates pathways enriched in hypomethylated genes. Gene rank plots show position of genes being part of a pathway set within the ordering of all genes based on methylation difference in mCG. Pathways with p value larger than 0.05 shown in red.

**Supplementary Figure 10.**
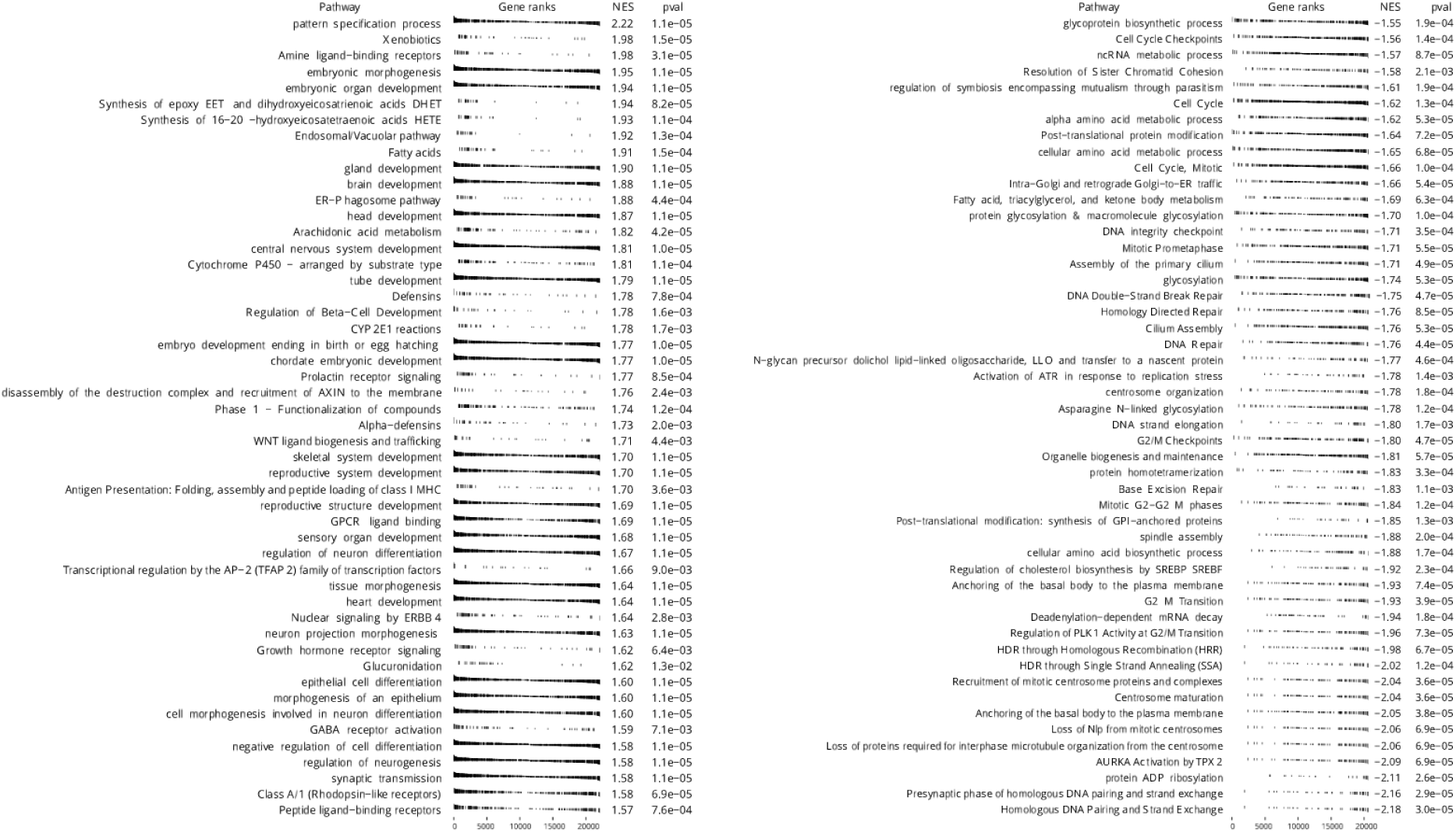
Top 50 enriched pathways for genes differentially methylated in CH context between in vitro neurons and in vivo neurons. Pathways were ranked by enrichment score (NES), whereas positive NES indicates pathways enriched in genes hypermethylated for CH in in vitro neurons, while negative NES indicates pathways enriched in hypomethylated genes. Gene rank plots shows the position of genes being part of a pathway set within the ordering of all genes based on methylation difference in mCH.

**Supplementary Figure 11.**
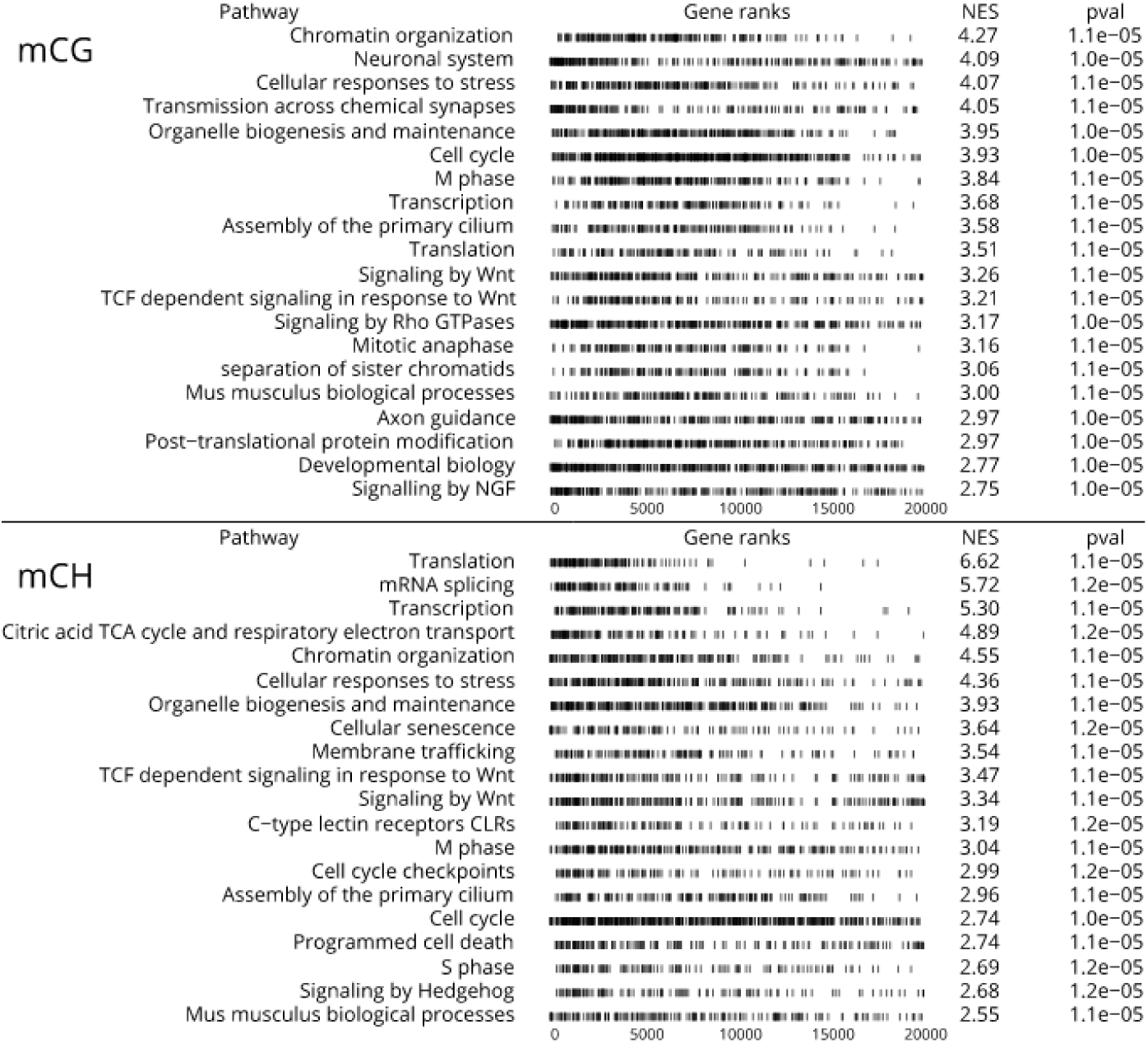
Enriched pathways for genes with similar methylation patterns for in vitro neurons and in vivo neurons. Pathways were ranked by enrichment score (NES), based on similarity between in vitro and in vivo neurons and dissimilarity to glia and fetal brain in mCG and mCH context respectively. Gene rank plots show the position of genes being part of a pathway set within the ranking of all genes.

